# Brainstem fMRI signaling of surprise across different types of deviant stimuli

**DOI:** 10.1101/2022.07.25.501374

**Authors:** Audrey Mazancieux, Franck Mauconduit, Alexis Amadon, Jan Willem de Gee, Tobias Donner, Florent Meyniel

## Abstract

The detection of deviant stimuli is crucial to orient and adapt our behavior. Previous work showed that infrequent (hence deviant) stimuli elicit phasic activation of the brainstem locus coeruleus (LC), which releases noradrenaline and controls central arousal. However, it is unclear whether the detection of behaviorally-relevant deviant events selectively trigger LC responses, or also other neuromodulatory systems related to dopamine, acetylcholine, and serotonin. Here, we combined human fMRI recordings optimized for brainstem imaging with pupillometry (a peripheral marker of central arousal) to perform a mapping of deviant-related responses in subcortical structures. Participants had to detect deviant items in a “local-global” paradigm that distinguishes between deviance based on the stimulus probability and the sequence structure. fMRI responses to deviant stimuli were quite distributed, detected in the LC but also other subcortical nuclei and many cortical areas. Both types of deviance elicited responses in the pupil, LC and other neuromodulatory systems. Our results reveal that the detection of task-relevant deviant items recruits the same multiple subcortical systems across computationally different types of deviance.

## Introduction

Detecting deviant stimuli (i.e. stimuli that violate some regularity) is crucial in a variety of processes, such as learning under uncertainty (Soltani and Izquierdo 2019), interacting in a flexible manner with the environment (Devauges and Sara 1990; Janitzky et al. 2015), and orienting behavior (Susan J. Sara and Bouret 2012). In terms of mechanisms, it seems clearly established that deviance detection triggers the phasic, brain-wide release of noradrenaline from the locus coeruleus (LC) located in the brainstem (G. Aston-Jones et al. 1994; Hervé-Minvielle and Sara 1995; S. J. Sara, Vankov, and Hervé 1994; Poe et al. 2020; Susan J. Sara and Bouret 2012), especially when deviant items are behaviorally relevant and correctly detected (Rajkowski, Kubiak, and Aston-Jones 1994). This conclusion is supported mostly by studies in non-human animals, using electrophysiological recordings of LC neurons during oddball tasks, in which frequent and rare stimuli are typically presented in a sequence: the LC responds specifically to the rare (hence deviant) stimulus (G. Aston-Jones et al. 1994; G. Aston-Jones, Rajkowski, and Kubiak 1997; Rajkowski, Kubiak, and Aston-Jones 1994; Foote, Aston-Jones, and Bloom 1980; Swick et al. 1994). Studies in humans provided converging evidence: the fMRI signal in the LC region increased after deviant stimuli in oddball tasks (Krebs et al. 2018; Murphy et al. 2014).

However, this body of work leaves unclear the anatomical specificity of deviant-related responses : Are they specific to the LC or shared across multiple other subcortical structures, notably neuromodulatory centers? The latter seems likely because deviance detection overlaps with other notions like novelty (Debener et al. 2005; Schomaker and Meeter 2015) and unexpectedness (Reichardt, Polner, and Simor 2020; Verleger and Śmigasiewicz 2016) when deviance is defined by rareness, and salience (Harsay et al. 2012) are known to implicate noradrenaline but also other neuromodulators (Bassareo, De Luca, and Di Chiara 2002; Brown et al. 2015; Bunzeck and Düzel 2006; Caldenhove et al. 2017; Glimcher 2011; Pessiglione et al. 2006; Ungless 2004; Vazey, Moorman, and Aston-Jones 2018; Kutlu et al. 2021). For instance, dopamine encodes unexpected stimuli in the form of reward prediction error (Glimcher 2011; Pessiglione et al. 2006) as well as salient stimuli related to novelty (Bassareo, De Luca, and Di Chiara 2002; Bunzeck and Düzel 2006; Kutlu et al. 2021).

Several pharmacological studies indicate that the deviance-related response recruits a large set of neuromodulatory systems. Propranolol (a blocker of the noradrenergic beta receptors) decreased fMRI signals in cortical regions that respond to deviant stimuli (Strange and Dolan 2007), but this effect is not specific to noradrenaline. For instance, the P300 event-related potential (ERP) was used as an indicator of LC activity supported by photoactivation studies in rats (Vazey et al., 2018), and it was found to be larger for deviant stimuli (Nieuwenhuis et al., 2005). However, this deviant-related P300 response was also found to be reduced following the administration of either scopolamine (a cholinergic antagonist) in rats (Ahnaou, Biermans, and Drinkenburg 2018) and humans (Caldenhove et al. 2017) or clonidine (a noradrenaline alpha receptor agonist) in humans (Brown et al. 2015).

Pupillometry has also often been used as an indirect marker of the LC activity and the existence of pupil responses to deviant stimuli is clearly established (Gilzenrat et al., 2010; Murphy et al., 2011; Nieuwenhuis et al., 2011; Preuschoff et al., 2011). However, a change in pupil size is not necessarily due to a change in LC activity (Megemont, McBurney-Lin, and Yang 2022) because other subcortical nuclei like the inferior colliculi (Joshi and Gold 2020; de Gee et al. 2017) and neuromodulators are at play such as acetylcholine from the basal forebrain (Reimer et al. 2016) and more indirectly serotonin from the raphe nucleus (Cazettes et al. 2021).

Here, we propose to measure deviant-related responses not only in the LC but also in other structures of the brainstem, notably in neuromodulatory centers. Direct, concurrent electrophysiological recording of multiple neuromodulatory centers is very difficult in non-human animals (Varazzani et al. 2015) and is not an option in humans. In contrast, fMRI can provide a complete coverage of the brainstem (and beyond), but brainstem fMRI is challenging due to the presence of larger physiological noise compared to cortex and to the very small size of the structures of interest, such as the LC (Liu et al. 2017). We thus used fMRI methods optimized for the brainstem (de Gee et al. 2017; Krebs et al. 2018) and delineated the LC (noradrenaline), the substantia nigra / ventral tegmental (SN/VTA; dopamine) and the superior and inferior colliculi (involved in pupil size and auditory processing (Ayala and Malmierca 2012; Malmierca et al. 2009; Joshi and Gold 2020)) based on the participant’s anatomy. Most fMRI studies that measured LC activity used anatomical atlases but this method is imprecise given its small size (Astafiev et al. 2010). We also included for comparison the activity of other neuromodulatory regions: the basal forebrain (BF) for acetylcholine, and the Raphe nucleus (RN) for serotonin (using atlases because they are more difficult to delineate individually) as well as other subcortical and cortical areas.

The studies mentioned so far used oddball (or similar) tasks (Foote, Aston-Jones, and Bloom 1980; G. Aston-Jones et al. 1994; G. Aston-Jones, Rajkowski, and Kubiak 1997; Rajkowski, Kubiak, and Aston-Jones 1994; Krebs et al. 2018; Murphy et al. 2014; Strange and Dolan 2007), in which deviant (oddball) items typically differ greatly from the standard items in terms of physical property (e.g. pitch difference) and probability, making them very salient. The deviant-related response may thus not reflect the detection of the deviant item *per se*, but downstream processes related to the salience or behavioral relevance of the deviant item. Here, we used the local-global paradigm (Bekinschtein et al. 2009) in which participants counted deviant items, making these deviant items behaviorally relevant (hence, salient). Interestingly, this paradigm differentiates between two types of deviant items: one based on the stimulus probability (just as in classical oddball tasks), which can be detected by simple mechanisms such as stimulus-specific adaptation (Ayala and Malmierca 2012; Malmierca et al. 2009); and another based on the structure of the sequence, which requires more elaborate processes to be detected (Dehaene et al. 2015) such as predictive coding (Heilbron and Chait 2018) and even awareness (King et al. 2013; Strauss et al. 2015). In other words, this paradigm dissociates the notion of task-relevant deviant items from a specific type of deviance. We investigated whether the responses to these task-relevant deviant items are similar or different between these two types of deviant items.

To anticipate our result we found that both types of deviant items elicit widespread fMRI responses in subcortical and cortical structures, which may correspond to the broadcasting of a task-related, salient event downstream of potentially different deviance detection systems.

## Results

### Distinguishing between different types of deviant items

The local-global paradigm presents sequences of stimuli that exhibit two nested levels of structure. At the local level, stimuli form patterns of 4 sounds with a fifth one that is either identical, forming a *locally standard* pattern xxxxx, or different, forming a *locally deviant* pattern xxxxY, see Figure 1A. The global level is characterized by the succession of patterns separated by short pauses: in each block of the task, one pattern is frequent (80% of patterns) while the other is rare (20%). The local and global properties, as well as sound identity were crossed in a full factorial, 2x2x2 design. In each block, participants were first familiarized with the frequent pattern only (which defines the block identity: xxxxx block or xxxxY block) and then presented with a few rare patterns (called global deviants) interleaved among the frequent ones. Participants listened to these patterns and were instructed to count the number of global deviant patterns. Twenty participants performed the task in the scanner where they had to count the number of rare patterns for a total of 4 sessions.

**Figure 1.**
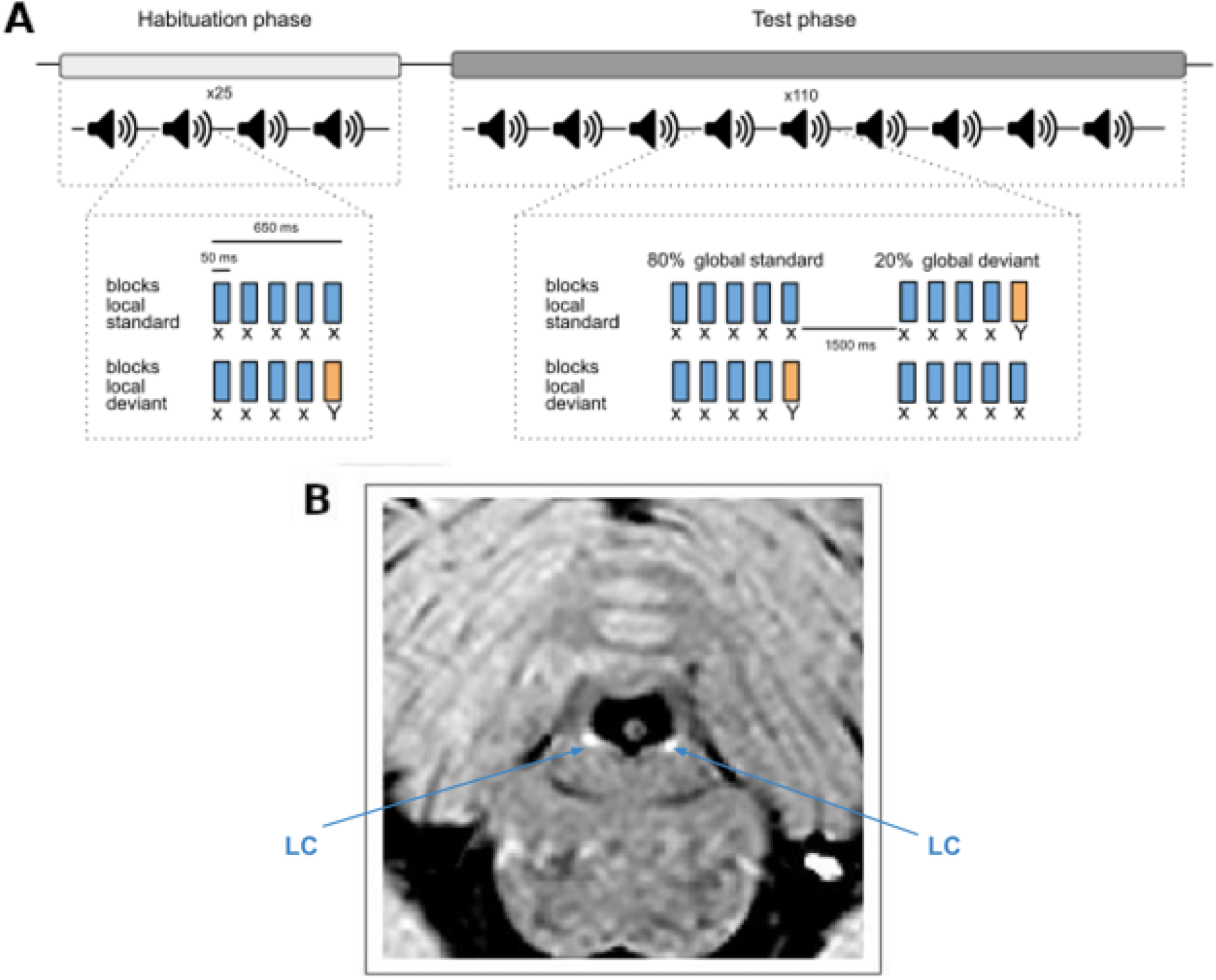
Task and example of anatomical images. **A)** The local-global paradigm. Patterns correspond either to 5 identical tones (“local standards” xxxxx) or 4 identical tones and a different one (“local deviants” xxxxY). During the habituation phase, only one of the two patterns is presented (called the global standard pattern). During the test phase, this pattern is presented 80% of the time, and the other pattern (called the global deviant pattern) is presented in 20% of the case. In total participants are presented with 4 different types of patterns: when the global standard (80%) is the local standard (xxxxx), then the global deviant (20%) is the local deviant (xxxxY); when the global standard (80%) is the local deviant (xxxxY), then the global deviant (20%) is the local standard (xxxxx). **B)** Example slice of the anatomical Turbo Spin Echo (TSE) image used to delineate the LC (appearing in hypersignal, i.e. brighter) in each participant.

Note that there is a key difference between the two types of global deviants. A rare xxxxY can be detected among frequent xxxxx based on low-level processes that operate on the probability of the sounds themselves in a sequence, like stimulus-specific adaptation, because the Y sound is extremely rare (occuring only with probability 0.2*⅕=0.04). In contrast, such low-level processes do not suffice to detect a rare xxxxx among frequent xxxxY because the x being the dominant sound (occurring with probability 0.8*1+0.2*⅘=0.96), xxxxY stands out more than xxxxx based on sound probability alone. A mechanism for the detection of a rare xxxxx must operate at a higher level, namely the sequence of sound patterns. The local-global paradigm thus allows a distinction between different computations for deviance detection, operating on the stimulus probability and sequence structure respectively, and possibly different mechanisms, such as bottom-up and top-down processes, see Discussion.

### Robust responses to global deviants in the pupil-linked arousal system

Pupil size is controlled by the autonomic nervous system. It provides a marker of arousal that is known to transiently increase when deviant stimuli are detected (Quirins et al. 2018). To characterize its response to global deviance, we performed a baseline-corrected, epoch-based analysis to isolate the phasic evoked response (see Methods). This analysis included only a subset of 13 participants who have a large-enough number of trials after artifact rejection (see Method). Pupil size exhibited a clear response to global deviants, with larger pupil size for rare patterns compared to frequent patterns (t_max_ = 6.85, p_max_ < 0.001, d_max_ = 1.90, cluster p_FWE_ < 0.001), see Figure 2A. Note that response was similar between the two types of global deviants (there was no significant effect of local deviance, t_max_ = 1.95, p_max_ = 0.075; or interaction between local and global deviances, t_max_ = 1.35, p_max_ = 0.201).

**Figure 2.**
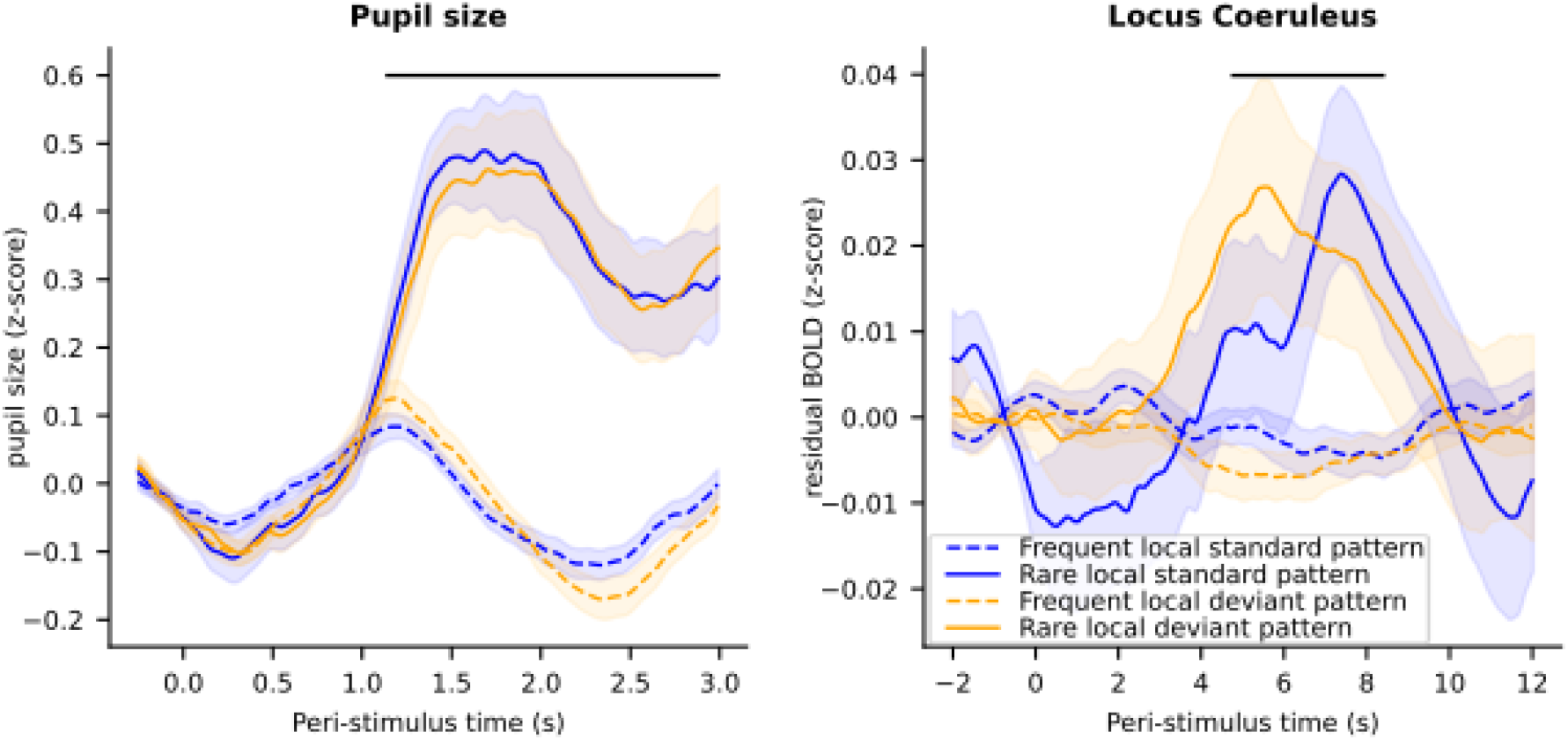
Epoch-based analyses of the 4 stimuli for pupil and L￼C data. **A) Pupil size** (z-score) evoked by the 4 types of patterns. B) Time course of fMRI activity (￼z-score) in the LC evoked by the 4 types of pattern. Error shading is standard error. Black dashed lines indicate significant clusters for the effect of rare patterns (global effect, p_FWE_<0.05).

### Global deviance transiently increases LC activity

Having established that both types of global deviants in the local-global paradigm elicit strong transient activation of the pupil-linked arousal system, we next investigated their effects in our primary region of interest (ROI), namely the LC. As for the pupil, we used epoch-based analyses. We extracted time series from the LC, removed several confounds (see methods), epoched the signal (from −2 to 12 s around the onset of patterns), and corrected for the baseline (subtracting the signal from −2 to 0 s). This analysis allowed us to track the fMRI activity of the LC in response to deviant stimuli with no assumption about the shape of the hemodynamic response, which may be different in subcortical structures compared to the cortex where the canonical hemodynamic responses has been defined (Friston et al. 2007). In addition, as for the pupil size analyses, the baseline correction removes the autocorrelation that may exist in the signal before and after the pattern onset, and captures the change in activity evoked by the pattern. Thus, this analysis captures the phasic activity of the LC and removes the tonic activity.

The LC activity showed a main effect of global deviance (t_max_ = 3.04, p_max_ = 0.006, d_max_ = 0.62, cluster p_FWE_ = 0.024) with a greater increase in fMRI activity for rare patterns compared to frequent patterns, see Figure 2B. No cluster was identified for either the local effect or for the interaction between the local and global deviance suggesting that responses were similar between the two types of global deviants.

### Mapping of deviance-related responses across regions of interest

We repeated the epoch-based analysis to quantify deviant-related responses in several other brain structures sorted into three categories. 1) The other neuromodulator nuclei included the SN/VTA (individually delineated, see Methods and Figure S5), the BF and the RN (based on anatomical atlases). 2) The other subcortical structures included the superior colliculi (involved in orienting responses), the inferior colliculi (involved in auditory processing), and the hippocampus (involved in sequence processing). 3) Cortical structures, where the fMRI signal-to-noise ratio is higher than in subcortical structures, included the superior temporal gyrus (previously identified to respond to rare patterns with the same task, see (Bekinschtein et al. 2009)), the primary auditory and visual cortices (corresponding to the calcarine and the superior temporal sulcus in Destrieux’s parcellation; Destrieux et al., 2010), and the rectus gyrus in the medial prefrontal cortex (which is part of the default mode network and thus not expected to respond to rare patterns, see section 1 of Supplementary Results for the effect of rare pattern in a full parcellation of the brain using the Destrieux’s parcellation).

Figure 3 shows the 4 types of stimuli for each ROI and Table 1 summarizes the corresponding statistics (see section 2 of Supplementary Results for a similar figure gathering the 2 types of rare and frequent patterns). All neuromodulator nuclei showed an increase in fMRI activity for rare patterns compared to frequent patterns (which remained significant when corrected for multiple comparisons across time, except in the BF). The dopaminergic SN/VTA is the structure that elicits the largest response. For the LC, the maximum effect size was smaller than in the SN/VTA and the RN, probably due to the smaller size of the LC ROI. The ventral medial prefrontal cortex and the hippocampus showed the reverse pattern later in the time window: fMRI activity in these regions was higher for frequent stimuli compared to rare stimuli. The other structures (cortical and subcortical) all exhibited a larger response for rare patterns compared to frequent patterns. Larger responses were found in cortical ROIs than in sub-cortical areas, potentially due to their different sizes and signal-to-noise ratios. The shape of the response also differed between cortical and subcortical regions: cortical regions had a more canonical response with a clear peak while several subcortical regions exhibit a kind of plateau (see Figure 3). Note that caution should prevail when comparing these different structures. First, cortical and subcortical structure differ in terms of SNR. Second, within subcortical structures, 4 have been defined in native space and others using atlas render the latest less neuroanatomically accurate.

**Figure 3.**
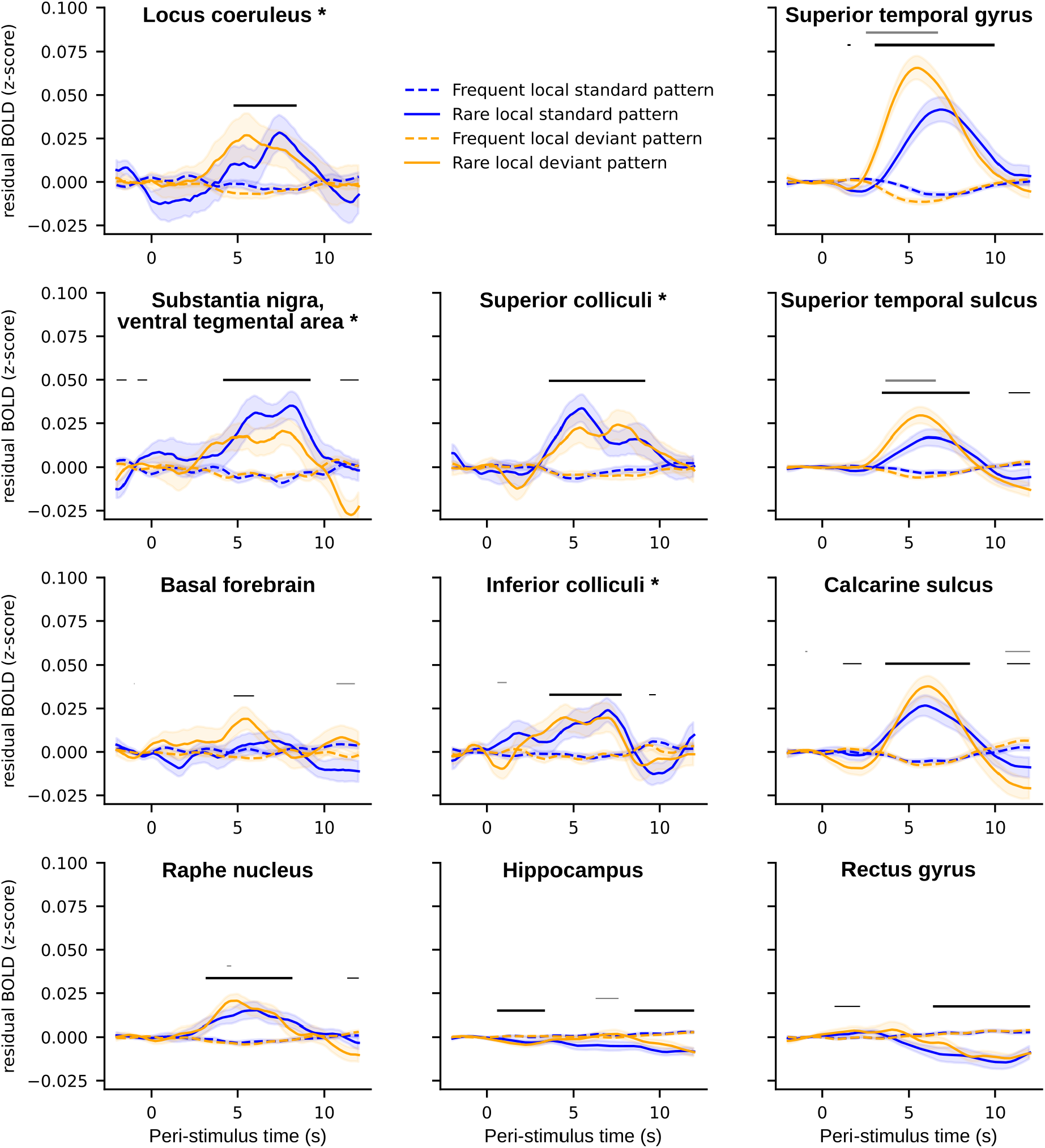
Time course of fMRI activity (z-score) evoked by the 4 types of patterns. The first column shows neuromodulator nuclei, the second column other subcortical ROIs, and the third column cortical ROIs. Stars indicate ROI defined in native space by manual delineation. Error shading is standard error. Bold black lines indicate significant clusters for the effect of rare patterns (global effect) and bold gray lines indicate clusters for the interaction between the global effect and the local effect (p_FWE_<0.05). Non-bold lines indicate time points with significant differences (p < 0.05) but cluster p_FWE_>0.05).

**Table 1.** Statistics for the global effect (max t-values, max p-values, max Cohen’s d, and p-values for clusters with FWE correction) for all ROIs.

| **Neuromodulation-related nuclei** | **Other subcortical nuclei** | **Cortical areas** |
| --- | --- | --- |
| **LC**<br>$t_{\max} = 3.04$<br>$p_{\max} = 0.006$<br>$d_{\max} = 0.62$<br>cluster $p_{\text{FWE}} = 0.023$ | **Superior colliculi**<br>$t_{\max} = 4.24$<br>$p_{\max} < 0.001$<br>$d_{\max} = 0.86$<br>cluster $p_{\text{FWE}} = 0.005$ | **Superior temporal gyrus**<br>$t_{\max} = 8.52$<br>$p_{\max} < 0.001$<br>$d_{\max} = 1.74$<br>cluster $p_{\text{FWE}} < 0.001$ |
| **SN/VTA**<br>$t_{\max} = 5.52$<br>$p_{\max} < 0.001$<br>$d_{\max} = 1.13$<br>cluster $p_{\text{FWE}} < 0.001$ | **Inferior colliculi**<br>$t_{\max} = 3.46$<br>$p_{\max} = 0.002$<br>$d_{\max} = 0.71$<br>cluster $p_{\text{FWE}} = 0.004$ | **Calcarine sulcus**<br>$t_{\max} = 6.31$<br>$p_{\max} < 0.001$<br>$d_{\max} = 1.29$<br>cluster $p_{\text{FWE}} < 0.001$ |
| **BF**<br>$t_{\max} = 2.35$<br>$p_{\max} = 0.027$<br>$d_{\max} = 0.48$<br>cluster $p_{\text{FWE}} = 0.159$ | **Hippocampus**<br>$t_{\max} = -4.69$<br>$p_{\max} < 0.001$<br>$d_{\max} = -0.96$<br>cluster $p_{\text{FWE}} = 0.022$ | **Superior temporal sulcus**<br>$t_{\max} = 5.53$<br>$p_{\max} < 0.001$<br>$d_{\max} = 1.12$<br>cluster $p_{\text{FWE}} < 0.001$ |
| **RN**<br>$t_{\max} = 4.70$<br>$p_{\max} < 0.001$<br>$d_{\max} = 0.96$<br>cluster $p_{\text{FWE}} = 0.002$ | | **Rectus gyrus**<br>$t_{\max} = -5.05$<br>$p_{\max} < 0.001$<br>$d_{\max} = -1.03$<br>cluster $p_{\text{FWE}} = 0.006$ |

No ROI showed a main effect for local deviance. Interaction between the local and global effect was significant only for cortical areas involved in auditory processing, namely the superior temporal gyrus (t_max_ = 6.08, p_max_ < 0.001, d_max_ = 1.24, cluster p_FWE_ < 0.001) and the superior temporal sulcus (t_max_ = 3.84, p_max_ < 0.001, d_max_ = 0.78, cluster p_FWE_ = 0.045). In these regions, the signal time courses (Figure 3) indicate that the interaction originates from the fact that the rare xxxxY patterns (the stimulus-probability deviant) elicited a higher and earlier response than the rare xxxxx patterns (the structure deviant). To confirm this hypothesis, we performed a follow-up analysis to estimate the difference in response peaks between conditions. The superior temporal gyrus showed a significant difference in the peak of the response, which was significantly earlier for rare xxxxY than rare xxxxx patterns (estimated difference: 1.422 s, 95% confidence interval: 0.888, 2.004). A similar trend was observed in the superior temporal sulcus (mean: 0.672, 95% CI=[-0.102, 1.742]).

To rule out that the detection of global deviant-related responses depends on the specifics of our analysis approach, we compared frequent and rare patterns across brain structures using Finite Impulse Response (FIR) analyses and General Linear Model (GLM) analyses (see section 3 and 4 of Supplementary Results). Epoch-based analyses do not model the potential superposition of effects of the current and previous patterns in the time window of interest. In contrast, FIR and GLM analyses are designed to model this superposition, and differ in their assumptions about the hemodynamic response (which is unconstrained or assumed to be canonical, respectively). The epoch-based analysis also contains a baseline-correction that aims to suppress endogenous fluctuations in the signal - therefore focusing on phasic activity - which are ignored by the FIR and GLM analysis. Those three analyses are thus complementary.

Overall, results were qualitatively consistent across the three types of analyses suggesting that the global effect does not depend on the type of analyses we performed. The only notable qualitative differences concern the hippocampus and the ventral medial prefrontal cortex where the late negative effect of global deviance found in epoch-based analyses (and FIR analyses) differed from the positive (non-significant) effect was found with GLM analyses, probably because the late difference is not well captured by the canonical response function.

### Anatomical specificity of the response to global deviants around the in the LC region

The global deviant-related response is very much distributed in cortical and subcortical structures, raising the concern that the effect found in the LC may not be specific to this region but instead widespread within the pons. To test for the anatomical specificity of global deviance within the pons, we repeated the same epoch-based analyses but after shifting the ROI corresponding to the LC in space. In the native space of each participants, we shifted this ROI toward the from of the head (from 1 to 5 voxels, i.e. +2 to +10 mm leading to 5 new ROIs, see Figure 4B) and toward the back of the head, which falls in the fourth ventricle (from 1 to 3 voxels, i.e. −2 to −6 mm, leading to 3 new ROIs, see Figure 4B). This axis is more relevant for the shift of the LC ROI than shifting toward left or right because the LC is a bilateral structure, and more relevant than toward the spinal cord or the midbrain because the LC is elongated along this axis. For each shift of the ROI we extracted the corresponding time-serie and performed epoch-based analysis for global deviant and global standards stimuli.

**Figure 4.**
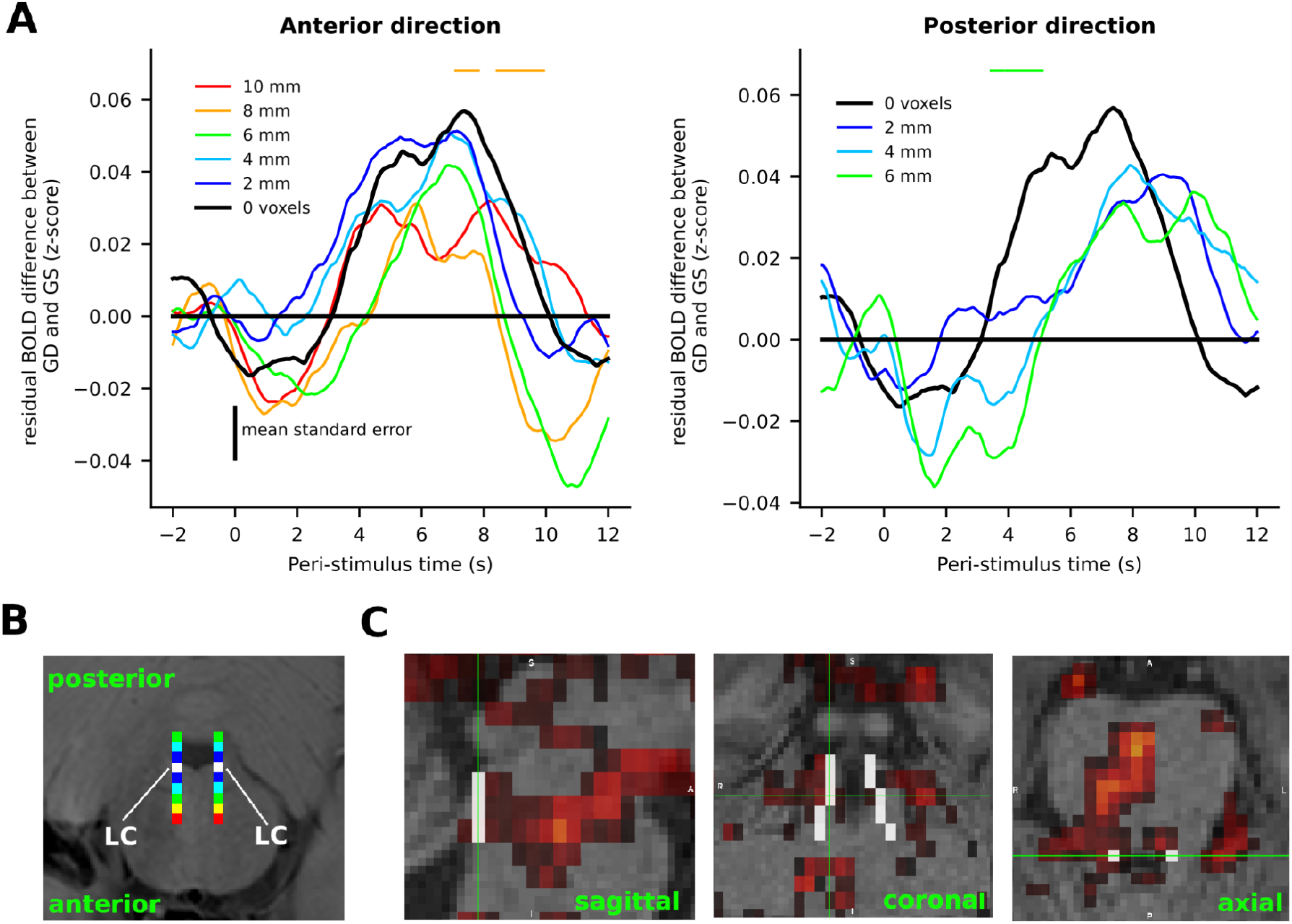
Anatomical specificity around the LC in the pons. A) Time course of the effect of global deviance (difference in z-scored fMRI activity between rare and freque￼nt patterns) for different shifts of the anatomically-defined LC ROI in millimeters (2 mm corresponds to 1 voxel). Left panel refers to shifts toward the anterior direction of the pons. Right panel refers to shifts toward the posterior direction of the pons. Black line refers to the original, non-shifted LC ROI. Colored horizontal dashed lines refer to identified clusters for the difference between the corresponding color and the black line. None of them remains significant after correction for multiple comparisons (one-sided tests). B) Example location of the LC in the native space of one participant (in white) and the same ROI shifted along the antero-posterior axis (gradients from blue to red). C) Statistical z-map for the effect of global deviant stimuli, thresholded at z = 2.8 (corresponding to p = 0.005). White voxels correspond to the LC atlas from Keren et al. (2015).

**Figure 5.**
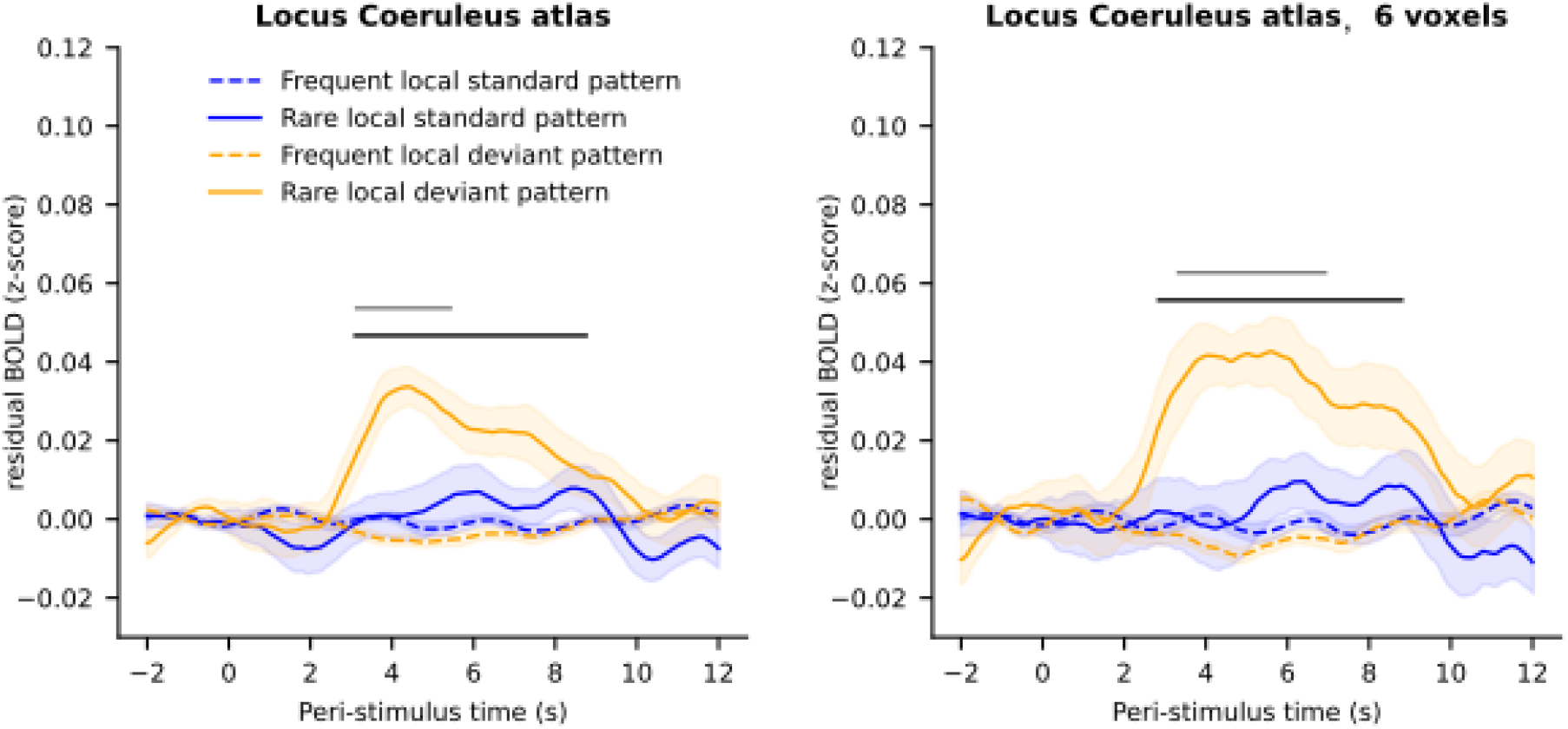
Anatomical specificity assessed with a normalized atlas of the LC. Time course of fMRI activity (z-score) in the LC evoked by the 4 types of patterns based on an anatomical atlas (left) and a selection of 6 voxels in the most anterior part of this atlas (right). Error shading is standard error. Black and gray dashed lines indicate significant clusters for the global effect and the interaction between global and local effects, respectively (p_FWE_<0.05).

Figure 4A shows the time course of the difference in fMRI signals between rare patterns and frequent patterns, for different shifts of the LC ROI (see section 5 of Supplementary Results for the same analysis for the xxxxY pattern and the rare xxxxx pattern separately). The effect of global deviance overall decreased as larger shifts were applied: the maximum difference between rare and frequent pattern changed from 0.057 (sd = 0.020) originally to 0.051 (sd = 0.020), 0.051 (sd = 0.022), 0.042 (sd = 0.021), 0.031 (sd = 0.019), and 0.032 (sd = 0.020) in the anterior direction (+2 mm to +10 mm shifts) and to 0.041 (sd = 0.022), 0.043 (sd = 0.018), 0.036 (sd = 0.020) in the posterior direction (−2 mm to −6 mm shift). Only the signal for the actual LC (no shift) and the signal for the +2 mm and −4 mm shift showed a significant effect of global deviance (no shift: p_FWE_ = 0.023; +2 mm shift: p_FWE_ = 0.006; −4 mm shift: p_FWE_ = 0.010). Direct comparisons of the global deviance in shifted and unshifted data showed time points with significant differences (p < 0.05 one-sided test, for shifts of +8 and −6 mm, but the corresponding cluster p_FWE_ remained > 0.05).

We also performed GLM, statistical mapping analysis for the effect of global deviant stimuli. Significant voxels in the brainstem showed some overlap with the LC atlas (see Figure 4C). Those results support an anatomical specificity of the effect of global deviance in the LC region compared to its vicinity, but without sharp boundaries.

### Comparison of LC activity in native space and atlas

The anatomical specificity of the effect of global deviance in the LC region can also be assessed by comparing the results obtained with anatomical delineation of the LC in each subject (in native space) to the expectedly less accurate ones obtained from a probabilistic atlas of the LC (in standardized space). We extracted fMRI time series based on an atlas of the LC (see Method section) that identified 10 voxels in standardized space. Note that our delineation in native space identified only a mean of 5.54 voxels for the LC (min = 4 voxels, max = 9 voxels, across participants). A comparison of the two approaches revealed that the voxels identified in native space were systematically in the more anterior part of the atlas of the LC, thus, closer to the midbrain, which is consistent with previous studies (Keren et al. 2009). Therefore, to allow a fair comparison with the individual delineations (Figure 2), we also performed a second analysis matched in voxel number (using 6 voxels in the atlas that were the most anterior). The two atlas-based analyses showed significant effects of global deviance (full atlas: t_max_ = 4.15, p_max_ < 0.001, d_max_ = 0.85, cluster p_FWE_ = 0.002; atlas with 6 voxels: t_max_ = 4.16, p_max_ < 0.001, d_max_ = 0.85, cluster p_FWE_ < 0.001). No significant cluster was identified for the effect of local deviance but the interaction between local deviance and global deviance was significant in both analyses (full atlas: t_max_ = 4.15, p_max_ < 0.001, d_max_ = 0.85, cluster p_FWE_ = 0.025; atlas with 6 voxels: t_max_ = 2.93, p_max_ = 0.009, d_max_ = 0.66, cluster p_FWE_ = 0.022). In contrast to the analysis performed after individual delineation of the LC, a region corresponding to the atlas exhibits a response to global deviance that is mostly driven by stimulus probability (rare xxxxY patterns). This is in line with the fact that the response to the rare xxxxY is largely shared by voxels in the vicinity of the individually, anatomically defined LC, whereas the response to the rare xxxxx patterns is not, but instead more specific to the LC itself (Figure S4A-B). If the atlas corresponds to voxels that are not exactly centered on the true, individually and anatomically defined LC of each participant, but instead in its vicinity, then the response to rare xxxxx patterns is expected to be reduced and even undetected.

### Functional connectivity of subcortical regions

To explore how the different subcortical structures relate to one another, we performed a correlation analysis of intrinsic signals among subcortical regions. We used the residual time series (after removing the linear effect of external common causes – the stimuli – and potential confounds: movement parameters, signal in the 4th ventricle, and physiological signals, see Figure 6A). The LC signals correlated significantly with the other neuromodulatory centers (RN, SN/VTA; with the notable exception of the BF), and with the IC that is known to be involved in auditory deviance detection (Ayala and Malmierca 2012). SN/VTA and the RN were the regions that correlated the most with other subcortical structures.

**Figure 6.**
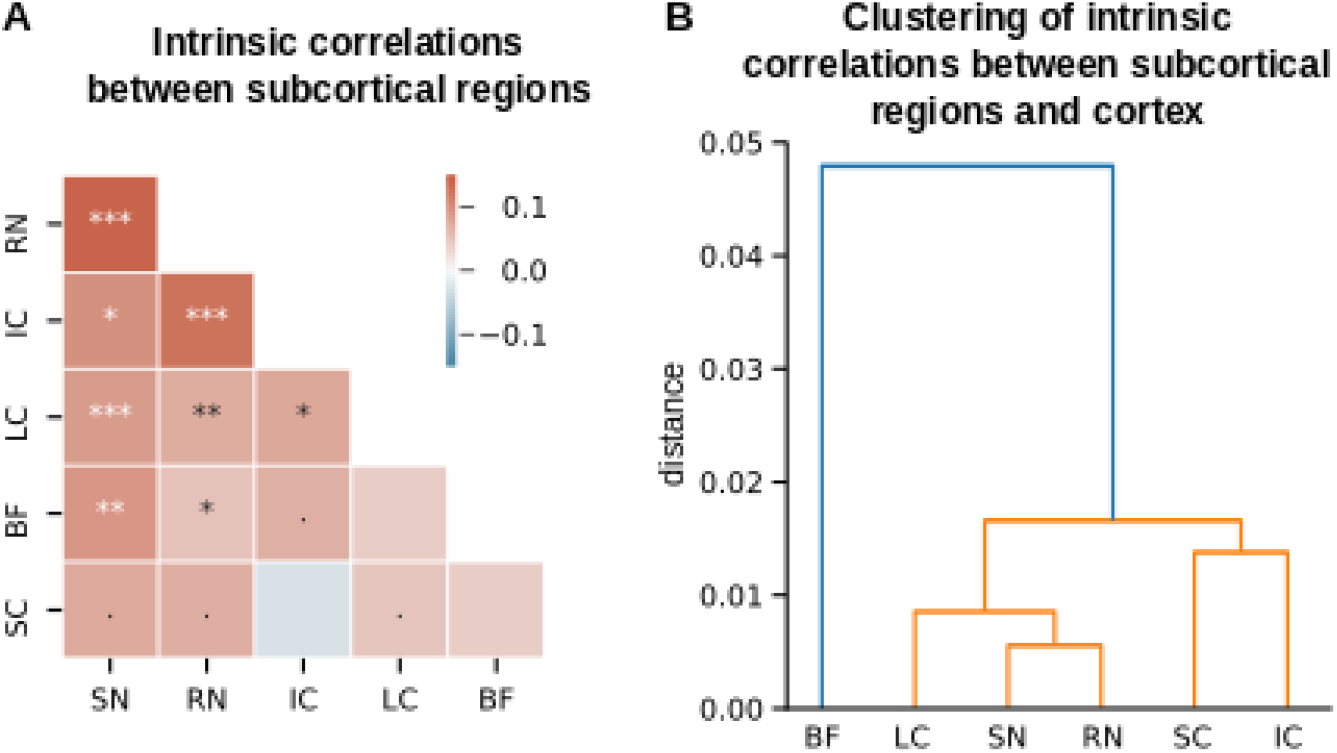
Functional connectivity of subcortical regions. (A) Correlation matrix of intrinsic signals across subcortical regions. Stars indicate Bonferroni-corrected significant correlations (*< 0.05, ** < 0.005, *** < 0.0005; the dot indicates p<0.05 uncorrected). (B) Regions were ordered based on a hierarchical clustering algorithm. Hierarchical clustering of subcortical regions based on their intrinsic correlations with cortical regions.

Correlations among subcortical structures are likely to be shaped by subcortical connections, and also afferent cortical connections. In a complementary analysis, we analyzed the pattern of correlations between cortical and subcortical signals. We measured the correlations of intrinsic signals between each subcortical structure and each cortical region that has substantial coverage in our field of view (>50% of the region, see Supplementary results). We then used the cortical-subcortical correlation matrix to cluster hierarchically the subcortical structures (Figure 6B). The cortical correlation profile of the BF appeared markedly different from the other subcortical regions and formed a separated cluster. Within the second cluster, the two regions of the tectal system (SC and IC), and the three remaining neuromodulatory centers (LC, SN, RN) were grouped together respectively. The subcortico-subcortical correlations (Figure 6) and the distance matrix between subcortical regions based on their cortical correlations (Figure 6) did not differ significantly between conditions (rare xxxxx blocs vs. rare xxxxY, all p > 0.14, except for the distance between the SC and the IC for which p = 0.03 without Bonferroni correction).

## Discussion

We performed a systematic response mapping in subcortical structures using fMRI coupled with pupillometry in a task that involves two types of deviants that requires computations based on the stimulus probability and sequence structure respectively. Global deviance evoked transient LC responses, which was our primary region of interest since it is well established that the central noradrenergic system vigorously responds to deviant stimuli (G. Aston-Jones et al. 1994; G. Aston-Jones, Rajkowski, and Kubiak 1997; Rajkowski, Kubiak, and Aston-Jones 1994). Similar responses were found in other neuromodulatory centers (the SN/VTA and the RN), other subcortical nuclei (the superior and the inferior colliculi), and cortical regions (the anterior and posterior superior temporal gyri, the primary auditory and visual cortices). Correlations of intrinsic signals revealed the functional similarity of the RN, SN/VTA and LC. Local and global deviances interacted in cortical responses related to auditory processing where global deviants elicited stronger and earlier responses when corresponding to a stimulus-probability deviant than a structure deviant. In contrast, subcortical structures (and the visual cortex) exhibited similar responses to both types of global deviants. This response to both types of global deviants showed some anatomical specificity to the LC region within the pons, because it decreased when moving away from the subject-specific, anatomically defined LC region, and became driven by stimulus-probability deviants when using a probabilistic, normalized atlas of the the LC. Pupil size exhibited similar responses to both types of global deviants.

The two types of deviant items in the local-global paradigm are by different brain circuits, according to previous electrophysiological studies. Local deviance (xxxxY vs. xxxxx) elicits an early response in sensory cortices whereas global deviance (rare vs. frequent patterns) elicit a later response that is distributed across brain areas and reaches the frontal lobe both in humans (Bekinschtein et al. 2009; Karoui et al. 2014; Wacongne et al. 2011; Dürschmid et al. 2016) and macaques (Chao et al. 2018). The effects of local deviance and global deviance are propagated across cortical areas through different frequency bands, the gamma band and beta-alpha band respectively (Karoui et al. 2014; Chao et al. 2018) which are distinct functional markers of bottom-up and top-down processes (Siegel, Donner, and Engel 2012; Wang 2010; Bastos et al. 2015). Our results focused on a comparison of the two types of global deviance and showed that rare patterns elicited stronger and earlier responses when they corresponded to the xxxxY patterns (stimulus probability deviant) than to the xxxxx pattern (structure deviant) in regions of the temporal lobe related to auditory processing, consistent with the idea that the detection of a rare xxxxx pattern recruits top-down processes (unfortunately, our brainstem-optimized partial brain coverage excluded most of the prefrontal cortex). We note that the distinction between global deviance based on stimulus probability and sequence structure is not tested in several previous studies (Bekinschtein et al. 2009; Karoui et al. 2014; Quirins et al. 2018), leaving unclear whether the global effect analyzed in those studies is driven by both types of global deviants, or just one.

In subcortical structures, in the pupil, and in the primary visual cortex, responses to both types of rare patterns were largely similar, in contrast to the temporal cortex. This similarity could indicate that they belong to a common final path, downstream of different circuits that detect different types of deviant items, that broadcasts the occurrence of a task-relevant, salient event. Subcortical structure such as the LC and the SN/VTA are known to be generally responsive to salient events, even when this salience is not due to the rareness of the event (Vazey, Moorman, and Aston-Jones 2018; Ungless 2004; Kutlu et al. 2021; Varazzani et al. 2015). Determining the source and target of neural activity in those cortico-subcortical networks would be valuable but would require better time-resolved techniques than fMRI, like electrophysiology, which is technically difficult to obtain. The LC in particular may play a central role in arousing the brain in response to task-relevant deviant stimuli. Studies in rodents and in primates showed that afferent LC inputs mainly come from subcortical nuclei and that in cortex, only the prefrontal region directly projects to the LC (Arnsten and Goldman-Rakic 1984; Luppi et al. 1995). Subcortical structures seem to be suitable candidates to signal the presence of rare xxxxY patterns, but as discussed above, they seem to lack the mechanisms to detect rare xxxxx patterns. The detection of the latter seems to rely on higher-order regions like the prefrontal cortex, which could directly signal those types to deviants to the LC.

In the local-global task, the increase in central arousal that follows rare patterns depends on the participants state of consciousness and their awareness of a sequence structure. Previous studies showed that the global deviance detection vanishes in patients with disorders of consciousness (Faugeras et al. 2012), when healthy subjects fall asleep (Strauss et al. 2015), and when they are not aware (or do not pay attention to) the task structure (Quirins et al. 2018). Interestingly, the effect of global deviance (notably rare xxxxx patterns) is more difficult to detect, and with a reduced extent, in brain recordings of macaque monkeys (Bellet et al. 2021; Chao et al. 2018; Uhrig, Dehaene, and Jarraya 2014; Jiang et al. 2022), for which global deviants are not behaviorally relevant and thus potentially not attended, compared to healthy human participants who are told about the existence of global deviants and often asked to count them (Bekinschtein et al. 2009; Karoui et al. 2014; Wacongne et al. 2011; Strauss et al. 2015; Quirins et al. 2018). Here, we also asked participants to count the global deviants, which probably enhanced their detection and the associated brain responses.

It is important to note that our design cannot distinguish between deviance detection *per se* (e.g.,. expectation violation signal) and the consequences of deviance detection (e.g., the identification of a task-relevant event). In addition, our fMRI data do not indicate where the deviance detection is computed in the brain for the two types of deviants. We can only speculate that expectation violation in some regions may serve to detect deviant items, possibly based on different circuits for the two deviant types and this detection may be communicated to (and enhanced by) some other regions due to its task-relevance.

LC responses are known to depend on attentional effects. During active oddball tasks, LC neurons exhibited a higher response when monkeys correctly (vs. incorrectly) detected rare stimuli (Rajkowski, Kubiak, and Aston-Jones 1994). More generally, there are state-dependent changes in tonic LC activity: higher tonic activity coincides with periods when monkeys have more motor activity that is irrelevant to the task; in contrast, periods of drowsiness, immobility, and eye-closure reduce LC activity (Rajkowski, Kubiak, and Aston-Jones 1994). Thus, phasic LC responses to deviant stimuli may occur for a particular level of tonic LC activity, when the subject is sufficiently focused (and not too much) on the current task (Gary Aston-Jones and Cohen 2005; Dubois et al. 2021). In this study, we reported baseline-corrected analyses (epoch-based analyses) to focus on the phasic component and suppress the additive effect of tonic fluctuations (but ignoring potential non-linear effects previously reported in spiking activity of LC neurons (Gary Aston-Jones and Cohen 2005) and in pupil size (Knapen et al. 2016)). In contrast, the non-baseline corrected analyses (see FIR and GLM analyses in Supplementary results) also focus on the phasic component but ignore the tonic fluctuations (the ones above 1/128 Hz that remain after preprocessing). We note that our results are consistent across baseline and non-baseline corrected analyses, which suggests in retrospect that the detection of rare patterns that manifests itself in an increased arousal of various brain structures was not masked by fast (above 1/128 Hz) fluctuations of tonic arousal levels (which would have penalized the non-baseline corrected analyses).

The current work will in addition be of methodological interest to people interested in the measure of LC activity with fMRI and more indirectly with pupillometry. The possibility to estimate the LC activity with fMRI is a contentious issue; doing so requires dedicated methods such as the subject-specific anatomical delineation of the LC, e.g. see technical comment (Astafiev et al. 2010; Eckert, Keren, and Aston-Jones 2010). We report a comparison of results obtained with a subject-specific delineation and a probabilistic, normalized anatomical atlas of the LC. Although both analyses showed an effect of global deviant patterns, this effect interacted with the local pattern type and was actually driven by the rare xxxxY pattern when using the atlas (there is no such interaction when using the subject-specific delineation, or in pupil size). Given what is observed in other brain regions, the pupil and previous work on the LC, we assume that the LC responds to both types of global deviants, and thus that the results obtained with subject-specific delineation are closer to the ground truth. In other words, using a subject-specific delineation (rather than an atlas) seems necessary in fMRI studies of the LC, despite being time and resource consuming. We also propose that the effect of the rare xxxxx patterns could be a quality check of a correct identification of the LC region, possibly in a trimmed down version of the local-global paradigm that only presents rare xxxxx among frequent xxxxY. A response to rare xxxxx patterns seems a more stringent test than a response to rare xxxxY pattern (as in oddball tasks) because we found responses to rare xxxxY patterns, but not rare xxxxx patterns, non-specifically across the pons (see Figure S4 in Supplementary Results).

Our results are also informative concerning the use of peripheral arousal (measured as non luminance-based change in pupil size) as an approximation of central arousal (more precisely, LC activity). The sensitivity of pupil size to LC activity is demonstrated based on direct LC recording in non-human animals (Reimer et al. 2016; Costa and Rudebeck 2016). However, those studies also demonstrated that this correlation is not specific to the LC activity, but also related to central acetylcholine (Reimer et al. 2016) and serotonin (Cazettes et al. 2021) levels. A consequence of this lack of specificity is that changes in pupil dilation may not reflect changes in LC activity (Megemont, McBurney-Lin, and Yang 2022). Here, we found an effect of the global deviance, without interaction with the local deviance, in both pupil size and fMRI activity in the LC region, suggesting that peripheral and central arousals are similar. Note that those similar effects could arise from the LC influencing pupil size (e.g. via the intermediolateral cell column, the Edinger-Westphal nucleus or the superior colliculi notably, (Joshi and Gold 2020)), or from a common input (e.g. the nucleus gigantocellularis that activates both the LC and the autonomic system (Susan J. Sara and Bouret 2012)). Those two hypotheses would have been supported by correlated responses to global deviance in the pupil and the LC region, but we did not find such a relationship, which was significant only between pupil size and fMRI activity in the SN/VTA region. This null result in the LC is not evidence for the absence of a relationship, notably because our analyses were limited by the small number of included trials and participants. The result found for the SN/VTA region (which replicates the one from (de Gee et al. 2017) in a different task) could simply be due to better data quality, this region being much larger than the LC (here, all effects were stronger in the SN/VTA than in the LC). The SN/VTA has no direct connection to the systems controlling pupil size (Joshi and Gold 2020) but it receives direct input from the LC (Susan J. Sara and Bouret 2012); the effect found in the SN/VTA could thus be due to an effect in the LC that our data failed to detect.

Overall, the current study showed deviant-related responses an effect of deviance that generalize across two types (stimulus probability and sequence structure) in many subcortical regions, including neuromodulatory centers, and several cortical regions. Our results are consistent with the idea that the detection of task-relevant deviant sound patterns triggers the arousal system through the activity of the LC. The LC likely gets inputs signaling the occurrence of a task-relevant event from higher regions (e.g. frontal areas) and in turn broadcasts surprise signals across the entire brain. Future work with better temporal resolution will need to determine the direction of neural signals between the interconnected neuromodulatory centers, other subcortical structures, and cortical areas that subtend a hierarchy of deviance mechanisms.

## Acknowledgments

This work was funded by the ANR #18-CE37-0010-01 CONFI-LEARN and by the ERC StG NEURAL-PROB #947105 to FM. The MRI facility was also partly supported by the European Union (FEDER-2007–2013, agreement #91–2015–004).

## Author contributions

F.M and A.M designed the MRI sequences. A.M. collected and analyzed the data. A.M, JW. dG, T.D, and F.M wrote the manuscript.

## Declaration of interests

There are no competing interests in this study.

## Materials and methods

### Participants

Twenty four participants (10 women) aged between 20 and 36 years (mean = 27.04, SD = 4.69) were enrolled in the experiment. This protocol was approved by a national ethics committee (Comité de Protection des Personnes Ile de France 3, approval #2018-A03195-50). Participants receive monetary compensation for their participation (80€ for 2 hours). They were right-handed based on self-report and had normal or corrected-to-normal vision.

### Stimuli and task

The task included 4 sessions of 10 minutes each and was run using Octave (version 6.1.0) and the Psychtoolbox (Brainard, 1997; Pelli, 1997; Kleiner et al, 2007) in the scanner. It was the same task as used in Bekinschtein et al. (2009). Stimuli are short auditory tones composed by 3 sinusoidal tones resulting in either a low-pitched sound (stimulus A composed by 350, 700, and 1400 Hz sinusoides) or a high-pitched sound (stimulus B: 500 Hz, 1000 Hz, and 2000 Hz sinusoides). Stimuli were presented in a sequence of patterns separated by pauses. A pattern consisted in four identical tones and a fifth that could be either the same (xxxxx; within-pattern standard, i.e. *local standard*) or different (xxxxY; within-pattern deviant, i.e. *local deviant*). The assignment of tones and patterns were counterbalanced across blocks (block of AAAAA and AAAAB vs. BBBBB and BBBBA). The duration of each tone was 50 ms and pattern duration was 650 ms with an inter-pattern interval of 1.500ms. During the habituation phase, participants were first exposed to only one pattern. During the test phase, participants were presented with either the same pattern as during habituation in 80% of the cases (*frequent pattern*) or with the other pattern in 20% of the cases (*rare pattern*). Figure 1A depicts a schematic representation of the task

Each session included 2 blocks in counterbalanced order: one where the habituation pattern was a local standard pattern (denoted xxxxx block) and one where it was the local deviant pattern (denoted xxxxY block). One block included 135 patterns (22 rare patterns and 113 frequent patterns including 25 ones for the habituation phase). During the task, participants had to listen to the pattern and count the number of rare patterns.

### MRI data collection and preprocessing

MRI data were acquired on a 3 Tesla scanner (Siemens, Prisma) with a 64-channel coil. In order to maximize the signal-to-noise ratio in LC, we acquired partial-brain functional echo planar images (EPI) images centered on the brainstem and oriented perpendicular to the floor of the fourth ventricle (and thus, main axis of the LC). We used the following parameters: TR = 1.25 s, TE = 30 ms, flip angle = 65°, 28 interleaved slices with a slice thickness of 3 mm and a multiband factor of 2. In-plane resolution was 2.0x2.0 mm. The encoding phase direction was from anterior to posterior. To estimate distortions, we acquired two volumes with opposite phase encoding directions. One volume was in the anterior to posterior direction (AP) and the other was in the other direction (PA), with TR = 4,800 ms, TE = 54 ms.

Two partial-brain Turbo Spin Echo (TSE) structural images, sometimes referred to as neuromelanin-sensitive (Chen et al., 2014; Sasaki et al., 2006) were acquired: one centered on the LC (e.g., DeGee et al., 2017; Keren et al., 2015) and others centered on the SN/VTA. Images were acquired with an in-plane resolution of 0.7x0.7 mm and reconstructed at 0.35x0.35 (TR = 675 ms, TE = 12 ms). We acquired 14 slices per TSE, slice thickness was 2 mm, oriented perpendicular to the floor of the fourth ventricle. We also acquired a whole-brain structural T1 image with an MPRAGE sequence for anatomical co-registration and the delineation of the IC and the SC with in-plane resolution of 1x1 mm and a slice thickness of 1 mm (TR = 2,300 ms, TE = 2.98 ms).

All preprocessing steps relied on SPM12 (Wellcome Trust Center for Neuroimaging, University College London) except the TOPUP correction that relied on FSL, using the python/FSL and python/SPM interfaces afforded by Nipype ( https://doi.org/10.5281/zenodo.596855). Slice-timing correction was referenced to the middle of each TR. Volumes were realigned onto the first volume of each session, and then onto the first volume of the first session. We also performed a TOPUP correction that estimates the susceptibility field using the AP/PA volumes and unwraps EPI images. Different coregistrations were made for different types of analyses. For those in native space analyses, EPI images were coregistered with the TSE images (either with the one centered on the LC to extract LC data, or the one centered on the SN/VTA to extract SN/VTA data) or with the T1 image (to extract IC and SC data). For normalized space analyses, the T1 image was first coregistered to the TSE image before normalization performed using the standard SPM template in the Montreal Neurological Institute (MNI) space.

### Physiological data collection and preprocessing

During the task, we recorded cardiac rhythm with a pulse oximeter and respiration with a belt. We modelled physiological signals using FSL PNM (Brooks et al., 2013) that creates physiological regressors for each slice of each volume. We selected estimates for the reference slice used in the slice-timing correction. We defined orders for each component as follows: 4 for the cardiac component, 3 for the respiratory component, and 1 for the interaction between the two. The total number of regressors modeling physiological signals was 18. One participant had no physiological recordings due to technical issues.

### Pupil size data collection and preprocessing

Pupil size was also recorded during scanning using an MRI-compatible EyeLink 1000 system. On raw data we performed the following preprocessing steps: (1) add a margin of 50 ms before and after the blinks detected by the EyeLink system, (2) interpolate the signal linearly within each blink, (3) low-pass filter (5 Hz) the data, (4) epoch the data within −0.5 to 3 s relative to each stimulus onset, (5) exclude epochs with a total blink duration exceeding 20% of the data. It is difficult to measure pupil size in the MRI scanner due to the distance between the eyes and the camera, the use of a mirror, and the partial occlusion by the antenna around the participant’s head. We excluded 11 participants for whom pupil size data was available on less than 20% of epochs. The number of participants included in the analyses related to pupil size was therefore 13.

### Definition of Regions of Interest (ROIs) and preprocessing

We delineated by hand for each participant ROIs in native space using the TSE images for the LC and SN/VTA (see Figure 1B for an example for one participant see section 8 of Supplementary Results for all participants), and the fourth ventricle, and the T1 image (for the IC and the SC). All masks were resampled to match the EPI resolution resulting in a probabilistic mask that was then transformed into a binary mask. Threshold probability of being part of the ROI was 0.05. We extracted time series from the EPI images using these masks. Anatomical landmark for the BF, the RN and to a lesser extent the hippocampus are less reliable in TSE and T1 images, thus, we used anatomical atlases in normalized space (maps from Zaborszky et al., 2008 for the BF; the Harvard Ascending Arousal Network atlas from Edlow et al., 2012 for the RN; the Harvard-Oxford cortical and subcortical structural atlases in FSL for the hippocampus). For comparisons between native and normalized space, we also used an anatomical atlas for the LC (Keren et al., 2015). Temporal signal-to-noise ratio in our main ROI - the native space LC - was 43.48 (sd = 5.46) (computed with the module TSNR from nipype, using a polynomial detrending of order 3).

For cortical ROIs, we used a complete parcellation of the whole brain (Destrieux et al., 2010) into 75 regions. For all these regions, we performed epoch-based analyses (see below) and reported the effect of global deviants in each region (see Supplementary results). From these regions, we selected the superior temporal gyrus and the superior temporal sulcus as the auditory processing regions, the calcarine for a primary visual processing region, and the gyrus rectus as a part of the default mode network. For each ROI, we preprocessed the signal by high-pass filtering (1/128 Hz).

### Epoch-based analyses of fMRI signals, baseline correction

We performed epoch-based analyses on fMRI time-series extracted from each ROI. We first linearly regressed out potential confounding variables (movement parameters, the time-series extracted from the fourth ventricle, and physiological regressors), and z-scored the residual signal per session. This signal was then upsampled (factor 1000, linear interpolation) and data was then epoched around each stimulus onset (time window: −2 s to 12) for each participant. Then, the baseline signal was subtracted from each epoch using a time window of −2 s to 0 s.

### Connectivity analysis and clustering

We estimated functional connectivity by calculating subject-level correlations on fMRI time-series extracted from each ROI across regions. We first linearly regressed out potential confounding variables (movement parameters, the time-series extracted from the fourth ventricle, and physiological regressors) as well as the effect of the stimuli, and z-scored the residual signal per session. We performed 2 types of connectivity analyses: one across subcortical regions (LC, SN/VTA, BF, RF, SC, and IC) and one between each of these regions and cortical regions with substantial coverage in our field of view (>50% of the region, see Supplementary results). We then performed hierarchical clustering based on those correlation matrices, using the module AgglomerativeClustering from scikit-learn (Pedregosa et al. 2012). The correlation matrix between subcortical structures was used as a distance metric (correlation distance), and the correlation matrix between subcortical and cortical structures was used as a feature matrix on which cosine distance was computed. The clustering algorithm used these distance matrices to cluster subcortical regions, using the average of distances as a criterion to merge clusters.

Finally, we repeated these 2 analyses for blocks with the rare xxxxx and blocks with the rare xxxxY. For each subject, we compared subcortico-subcortical correlations across blocks on the one hand, and cosine distance (between subcortico-cortical correlations) across blocks on the other hand. We performed a t-test at the group level to assess significance.

### Finite Impulse Response (FIR) analyses

FIR analyses model a number of successive post-stimulus time steps that allow to take into account stimuli that are presented to the participant during the time window of interest, controlling for potential superposition of effects. As for epoch-based analyses, the predefined time-window was from 0 sec to 12 sec around the onset of patterns and we added additional regressors (movement parameters, the fourth ventricle time-series, and physiological regressors) in our model. For these analyses, the fMRI signal was upsampled with a factor of 5. FIR analyses make no assumptions about the hemodynamic response. We only modelled the effect of rare patterns. At the group level, we tested whether the parameter estimates for these patterns differed from 0 by using a one sample cluster permutation test (cluster-forming and cluster-level alphas of 0.05, two-tailed tests, 10,000 permutations).

### Generalized Linear Model (GLM) analyses

As for FIR analyses, GLM-based analyses control for potential superposition of effects but assume the hemodynamic response to be canonical. One GLM was estimated on time-series per ROIs. The design matrix included the 4 types of patterns convolved with the canonical hemodynamic response function (as modeled in SPM) as well as additional regressors corresponding to movement parameters, time series in the fourth ventricle, and physiological regressors. Parameters (betas) were estimated at the subject level with an auto-regressive AR(1) model. We then computed the difference in parameter estimates between rare and frequent patterns, and tested for its significance (against 0) at the group-level using a t-test.

### Correction for multiple comparisons across time

As epoch-based analyses (of fMRI signals and pupil size) and FIR analyses require multiple comparisons across peri-stimulus times, family-wise error (FWE) correction for multiple comparisons was computed using a cluster-based permutation test (cluster-forming and cluster-level alphas of 0.05, two-tailed tests, 10,000 permutations) with the ‘mne’ package in Python (Gramfort et al., 2013).

## Supplementary results

### 1. Cortical effects of global deviance detection

Table S1 shows the group-level effect of global deviance for all anatomical parcels of the Destrieux parcellation (Destrieux et al. 2010). For each region (i.e. parcel), we report the maximum Cohen’s d for significant clusters after FWE correction (pFWE < 0.05).

**Table S1.** Maximum Cohen’s d for significant clusters after FWE correction (pFWE < 0.05) and coverage for each parcel (i.e. region) of the Destrieux parcellation. Bold font indicates the regions included in the functional connectivity analysis (i.e., more than 50% coverage). Blue regions are regions included in the manuscript.

| Code | Region | Coverage | Maximum Cohen'd |
| --- | --- | --- | --- |
| 1 | **Frontomarginal gyrus and sulcus** | **0.99** | **0** |
| 2 | **Inferior occipital gyrus and sulcus** | **1** | **0** |
| 3 | Paracentral lobule and sulcus | 0 | 0 |
| 4 | **Subcentral gyrus and sulcus** | **0.52** | **0.96** |
| 5 | **Transverse frontopolar gyrus and sulcus** | **0.58** | **0** |
| 6 | Anterior cingulate gyrus and sulcus | 0.05 | 0 |
| 7 | Middle-anterior cingulate gyrus and sulcus | 0 | 0 |
| 8 | Middle-posterior cingulate gyrus and sulcus | 0.01 | 0 |
| 9 | **Posterior-dorsal part of the cingulate gyrus** | **0.55** | **0** |
| 10 | **Posterior-ventral part of the cingulate gyrus** | **1** | **0** |
| 11 | **Cuneus** | **0.65** | **1.21** |
| 12 | Opercular part of the inferior frontal gyrus | 0.34 | 1.70 |
| 13 | Orbital part of the inferior frontal gyrus | 0.42 | **0.52** |
| 14 | Triangular part of the inferior frontal gyrus | 0.03 | 0.42 |
| 15 | Middle frontal gyrus | 0.06 | 0.57 |
| 16 | Superior frontal gyrus | 0.03 | 0 |
| 17 | **Long insular gyrus and central sulcus of the insula** | **0.98** | **0.65** |
| 18 | **Short insular gyrus** | **0.72** | **1.93** |
| 19 | **Middle occipital gyrus** | **0.75** | **0** |
| 20 | Superior occipital gyrus | 0.08 | 0 |
| 21 | Lateral occipito-temporal gyrus | 1 | 0 |
| 22 | **Lingual gyrus** | **1** | **0.97** |
| 23 | **Parahippocampal gyrus** | **1** | **0** |
| 24 | **Orbital gyrus** | **0.67** | **0.46** |
| 25 | **Angular gyrus** | **0.79** | **0** |
| 26 | **Supramarginal gyrus** | **0.55** | **1.14** |
| 27 | Superior parietal lobule | 0.06 | 1.24 |
| 28 | Postcentral gyrus | 0.001 | 0 |
| 29 | Precentral gyrus | 0.001 | 1.17 |
| 30 | **Precuneus** | **0.51** | **0** |
| 31 | **Rectus gyrus** | **0.72** | **0** |
| 32 | **Subcallosal gyrus** | **0.98** | **0.70** |
| 33 | **Anterior transverse temporal gyrus** | **1** | **1.30** |
| 34 | **Lateral aspect of the superior temporal gyrus** | **1** | **2.04** |
| 35 | **Planum polare of the superior temporal gyrus** | **1** | **1.21** |
| 36 | **Superior temporal gyrus** | **0.96** | **1.12** |
| 37 | **Inferior temporal gyrus** | **1** | **0.97** |
| 38 | **Middle temporal gyrus** | **1** | **0.45** |
| 39 | Horizontal ramus of the anterior segment of the lateral sulcus | 0.01 | 0.90 |
| **40** | **Vertical ramus of the anterior segment of the lateral sulcus** | **1** | **0** |
| **41** | **Posterior ramus of the lateral sulcus** | **0.95** | **0.84** |
| 42 | Occipital pole | 0.46 | 0.96 |
| **43** | **Temporal pole** | **1** | **0** |
| **44** | **Calcarine sulcus** | **1** | **1.29** |
| 45 | Central sulcus | 0.006 | 0.79 |
| 46 | Marginal branch (or part) of the cingulate sulcus | 0.04 | 0 |
| **47** | **Anterior circular insula sulcus** | **0.65** | **1.79** |
| **48** | **Inferior circular insula** | **1** | **1.31** |
| 49 | Superior segment of the circular sulcus of the insula | 0.19 | 1.36, |
| **50** | **Anterior transverse collateral sulcus** | **1** | **0** |
| **51** | **Posterior transverse collateral sulcus** | **1** | **0** |
| 52 | Inferior frontal sulcus | 0.15 | 0.68 |
| 53 | Middle frontal sulcus | 0.08 | 0.47 |
| 54 | Superior frontal sulcus | 0 | 0 |
| **55** | **Sulcus intermedius primus** | **1** | **1.02** |
| **56** | **Intraparietal sulcus and transverse parietal sulci** | **0.70** | **1.69** |
| **57** | **Middle occipital sulcus and lunatus sulcus** | **0.91** | **0.49** |
| **58** | **Superior occipital sulcus and transverse occipital sulcus** | **0.86** | **0.47** |
| **59** | **Anterior occipital sulcus** | **1** | **0** |
| **60** | **Lateral occipito-temporal sulcus** | **1** | **0** |
| **61** | **Medial occipito-temporal sulcus and lingual sulcus** | **1** | **0** |
| 62 | Lateral orbital sulcus | 0.01 | 0.61 |
| **63** | **Medial orbital sulcus** | **0.99** | **1.28** |
| **64** | **Orbital sulci (H-shaped sulci)** | **0.53** | **0.67** |
| **65** | **Parieto-occipital sulcus** | **0.96** | **0.64** |
| **66** | **Pericallosal sulcus** | **0.52** | **1.08** |
| 67 | Postcentral sulcus | 0.05 | 1.61 |
| 68 | Inferior precentral sulcus | 0.005 | 2.26 |
| 69 | Superior precentral sulcus | 0 | 0 |
| 70 | Suborbital sulcus | 0.38 | 0.53 |
| **71** | **Subparietal sulcus** | **0.72** | **0** |
| **72** | **Inferior temporal sulcus** | **1** | **0.91** |
| **73** | **Superior temporal sulcus** | **1** | **1.48** |
| **74** | **Transverse temporal sulcus** | **1** | **1.57** |

### 2. Effect of global deviance detection in all ROIs

Figure S1 shows the group-level effect of global deviance for each ROI (corresponding statistics can be found in Table 1 in the manuscript). All regions showed a significant (corrected for multiple comparisons across time points) increase in fMRI activity for rare patterns compared to frequent patterns (which remained significant when corrected for multiple comparisons across time) except for the BF, the hippocampus, and the rectus gyrus.

**Figure S1.**
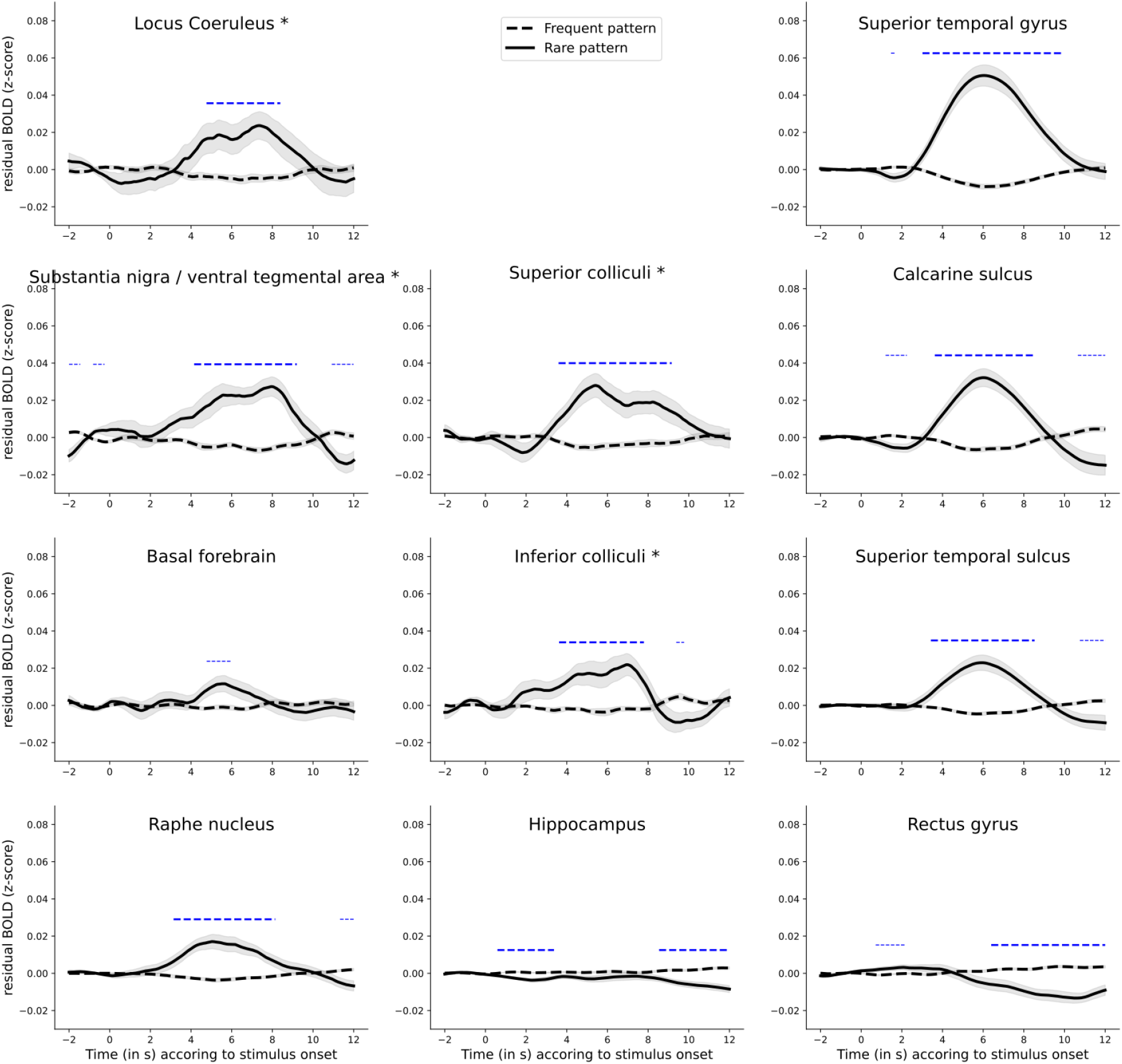
Time course of fMRI activity (z-score) evoked by rare patterns (bold line) and frequent patterns (dashed-line) in different brain structures. The first column shows neuromodulator nuclei, the second column other subcortical ROIs, and the third column cortical ROIs. Horizontal dashed blue lines correspond to clusters of significant difference (p < 0.05) and bold dashed blue lines to significant clusters after FWE correction (p_FWE_ < 0.05).

### 3. Effect of global deviance using FIR analyses

Figure S2 shows the group-level effect of global deviance for each ROI and Table S1 summarizes the corresponding statistics. As for epoch-based analyses, all neuromodulator nuclei showed an increase in fMRI activity for rare patterns. As for epoch-based analyses, only the hippocampus and the ventral medial prefrontal cortex showed no effect of global deviant pattern (no difference from 0). Note that for these analyses, the significant cluster identified for the LC did not reach significance (p_FWE_ = 0.067) when corrected for multiple comparisons across time.

**Table S2.** Statistics for the effect of rare patterns in FIR analyses (max t-values, max p-values, max Cohen’s d, and p-values for clusters with FWE correction), for all ROIs. When several clusters have been identified, the one with the higher max Cohen’s d is reported.

| **Neuromodulation-related nuclei** | **Other subcortical nuclei** | **Cortical areas** |
| --- | --- | --- |
| **LC**<br>$t_{\max} = 4.11$<br>$p_{\max} < 0.001$<br>$d_{\max} = 0.84$<br>cluster $p_{FWE} < 0.001$ | **Superior colliculi**<br>$t_{\max} = 4.19$<br>$p_{\max} < 0.001$<br>$d_{\max} = 0.86$<br>cluster $p_{FWE} = 0.002$ | **Superior temporal gyrus**<br>$t_{\max} = 8.86$<br>$p_{\max} = < 0.001$<br>$d_{\max} = 1.82$<br>cluster $p_{FWE} = < 0.001$ |
| **SN/VTA**<br>$t_{\max} = 6.28$<br>$p_{\max} < 0.001$<br>$d_{\max} = 1.28$<br>cluster $p_{FWE} < 0.001$ | **Inferior colliculi**<br>$t_{\max} = 4.36$<br>$p_{\max} < 0.001$<br>$d_{\max} = 0.89$<br>cluster $p_{FWE} < 0.001$ | **Calcarine sulcus**<br>$t_{\max} = 5.75$<br>$p_{\max} = < 0.001$<br>$d_{\max} = 1.17$<br>cluster $p_{FWE} = < 0.001$ |
| **BF**<br>$t_{\max} = 4.72$<br>$p_{\max} < 0.001$<br>$d_{\max} = 0.94$<br>cluster $p_{FWE} < 0.001$ | **Hippocampus**<br>$t_{\max} = 2.27$<br>$p_{\max} = 0.033$<br>$d_{\max} = 0.46$<br>cluster $p_{FWE} = 0.277$ | **Superior temporal sulcus**<br>$t_{\max} = 5.74$<br>$p_{\max} = < 0.001$<br>$d_{\max} = 1.17$<br>cluster $p_{FWE} = < 0.001$ |
| **RN**<br>$t_{\max} = 5.03$<br>$p_{\max} < 0.001$<br>$d_{\max} = 1.03$<br>cluster $p_{FWE} < 0.001$ | | **Rectus gyrus**<br>$t_{\max} = -7.83$<br>$p_{\max} = < 0.001$<br>$d_{\max} = -1.60$<br>cluster $p_{FWE} = < 0.001$ |

**Figure S2.**
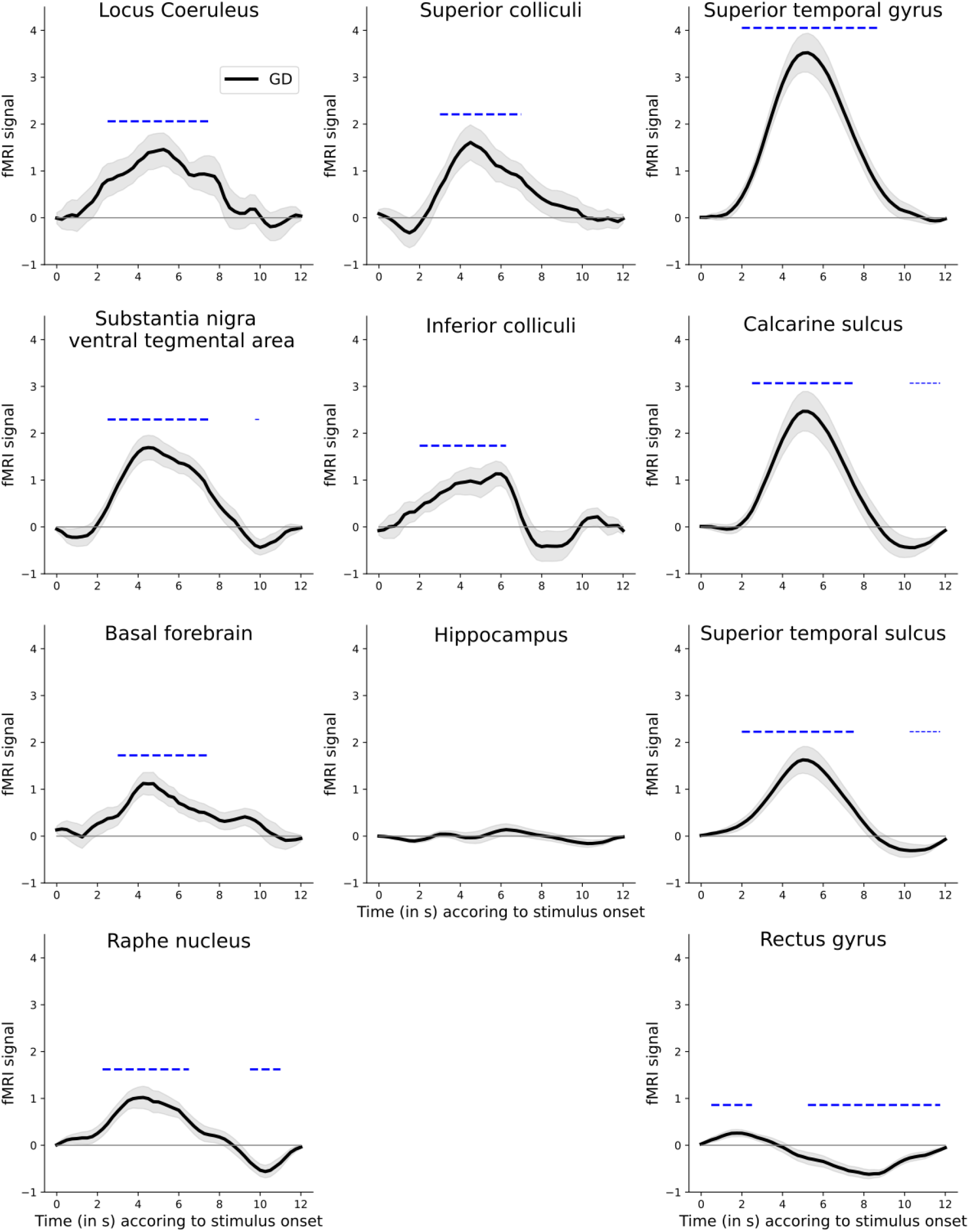
Time course of fMRI activity estimated using a FIR model for rare patterns. The first column shows neuromodulator nuclei, the second column other subcortical ROIs, and the third column cortical ROIs. Error shading is standard error. Blue dashed lines indicate significant clusters different from 0 (p_FWE_<0.05).

### 4. Effect of global deviance using GLM analyses

Figure S3 shows the distribution of single-subject estimates for the effect of global deviance for each ROI and Table S2 summarizes the corresponding statistics. All regions showed an increase in fMRI activity for rare patterns compared to frequent patterns.

**Table S3.**
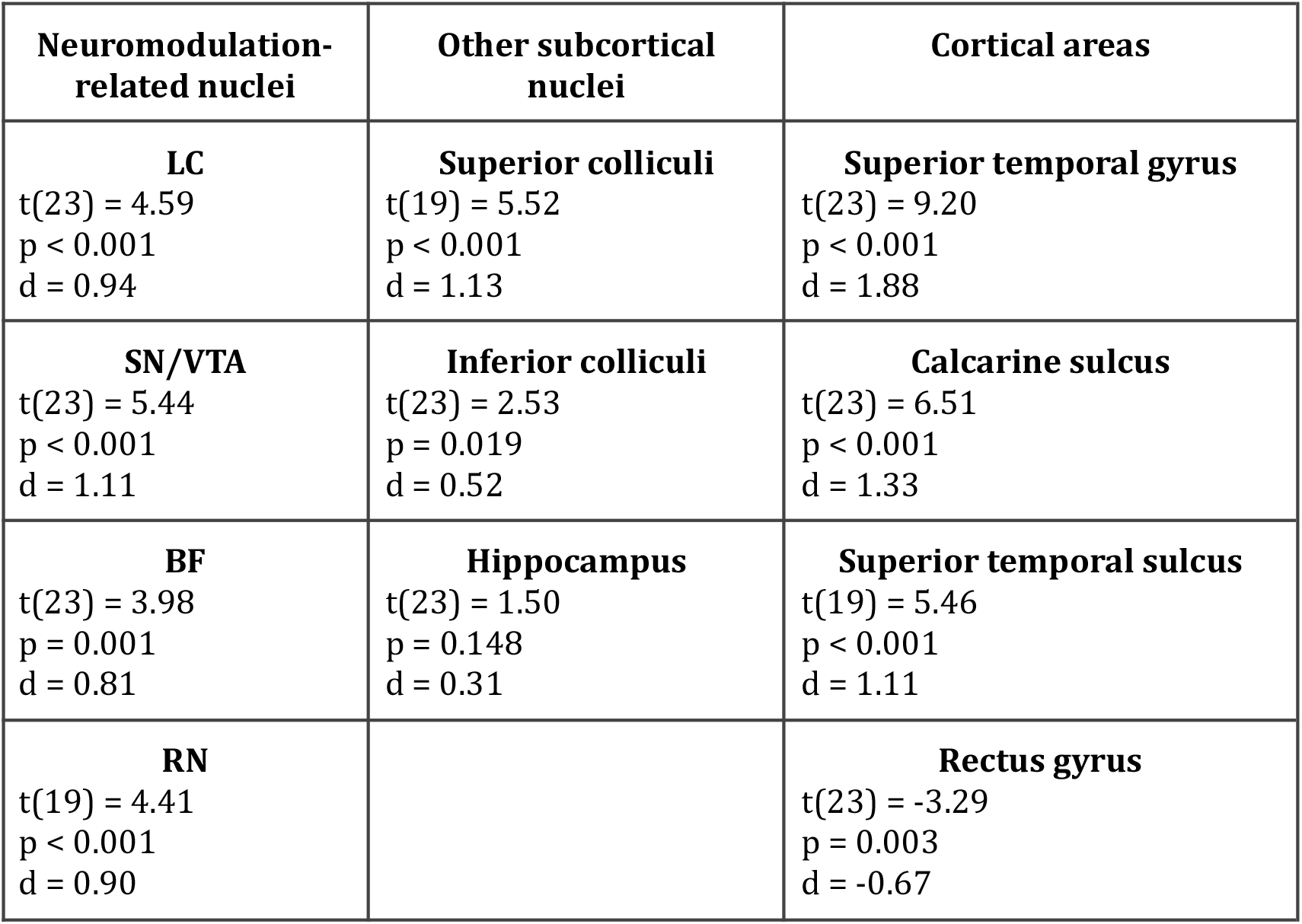
Statistics for the difference between the effects of rare and frequent patterns in GLM analyses (t-values, p-values, and Cohen’s d), for all ROIs.

**Figure S3.**
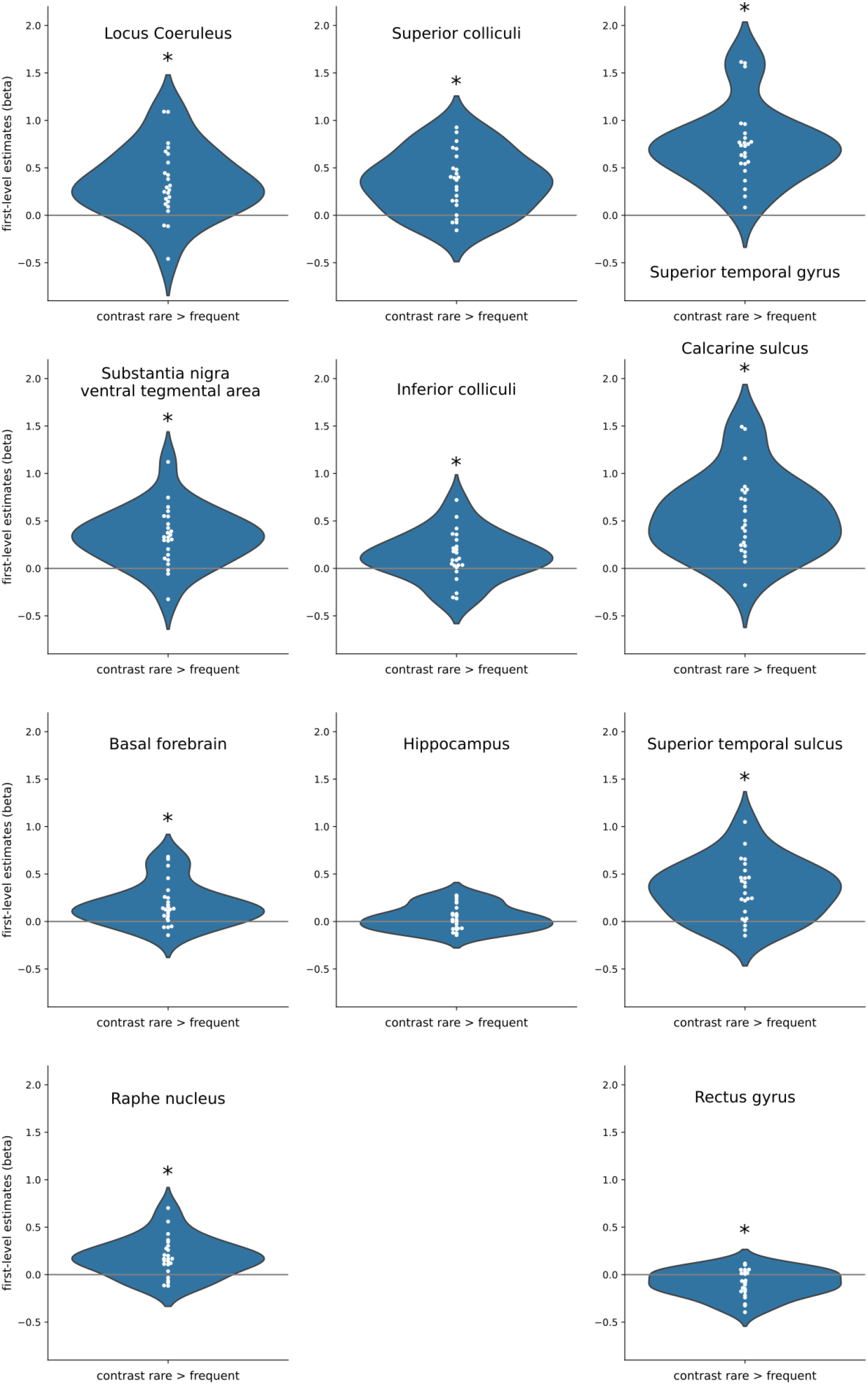
Single-subject estimates for the effect of rare patterns in different brain structures. The first column shows neuromodulator nuclei, the second column other subcortical ROIs, and the third column cortical ROIs. Stars indicate group-level significance (p < 0.05).

### 5. Anatomical specificity of the response each rare pattern around the in the LC region

The analyses that compared the LC defined anatomically on a subject-basis (in native space) and from an atlas (normalized space) led to different results, notably for the rare xxxxx stimulus that elicited responses in the case of subject-specific delineation but no response in the case of the atlas. By contrast, the rare xxxxY patterns elicited a response in both cases, and it was generally stronger than for rare xxxxx, notably in cortex. The atlas is likely to correspond only imprecisely to the LC of each individual, because of between-subject anatomical variability in the shape of the LC itself and of the brain at large, which undermines the normalization process (see Liu et al. 2017; de Gee et al. 2017). The different results between the subject-specific delineation and the atlas suggest that, unlike the response to rare xxxxY, the response to rare xxxxx patterns could be confined to the LC itself compared to its surroundings. To test this hypothesis, we repeated the anatomical specificity analysis (Figure 4) but for the two types of global deviant.

Figure S4A shows that overall, the magnitude of responses to the rare xxxxY remains the same when shifting the LC ROI away. The difference between the original ROI and the shifted ROI even shows a higher significant response when shifting by 2 mm and 4 mm in the anterior direction, and 6 mm in the posterior direction (cluster-level pFWE < 0.05, cluster-forming threshold: p<0.05). By contrast, Figure S4B shows that overall, the magnitude of responses to the rare xxxxx quickly faded when shifting the LC ROI away; the difference between the original ROI and the shifted ROI reached significance when shifting by 8 mm in the anterior direction (cluster-level pFWE < 0.05, cluster-forming threshold: p<0.05).

**Figure S4.**
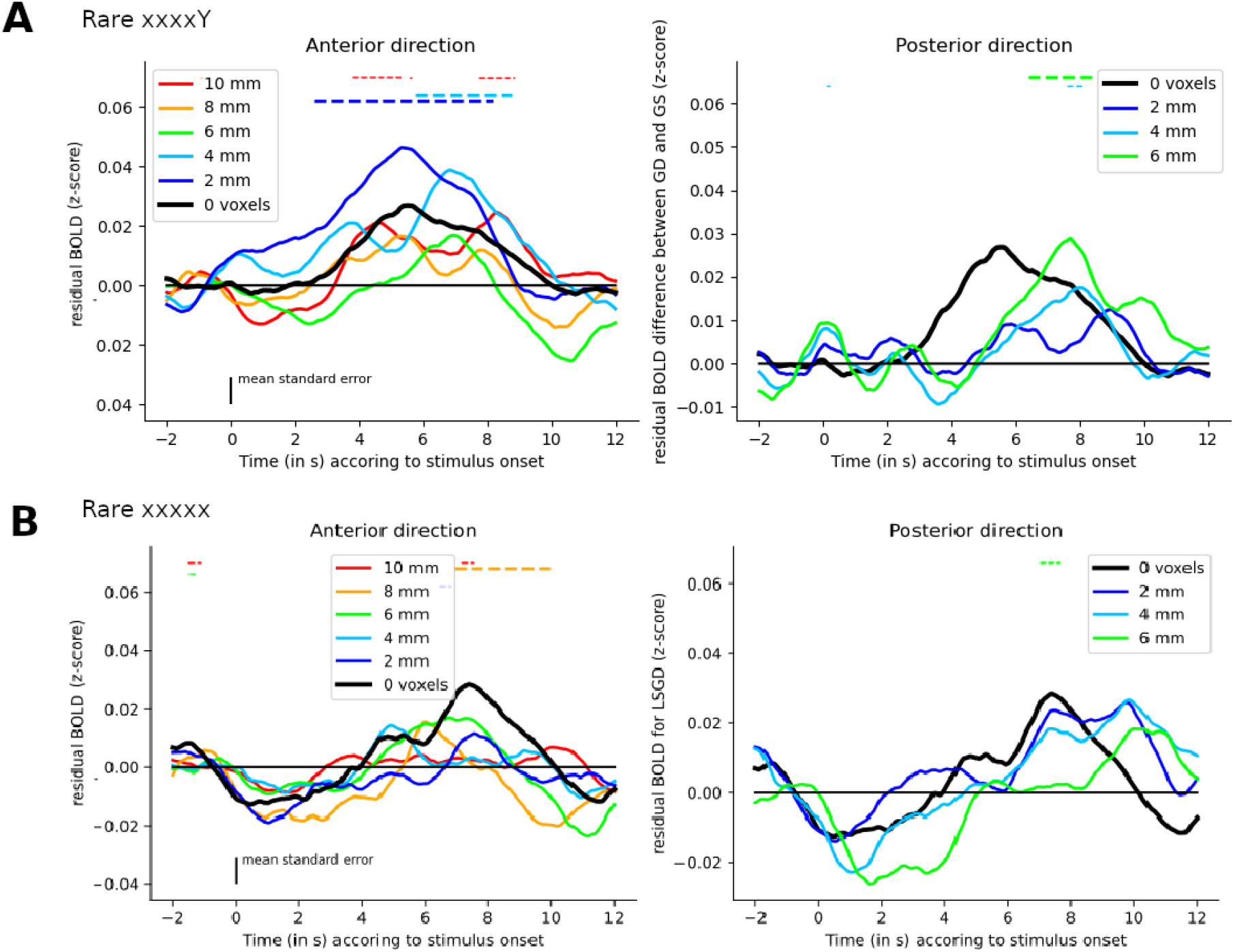
Time course of the z-scored fMRI activity for the two types of deviants or different shifts of the anatomically-defined LC ROI in millimeters. Two mm corresponds to 1 voxel. Colored horizontal dashed lines indicate the significant clusters (p<0.05; FWE corrected) identified for the difference between the corresponding color and the black line. A) For the rare xxxxY stimulus. Left panel refers to shifts toward the front of the head. Right panel refers to shifts toward the back of the head (see Figure 4B). Black line refers to the original, non-shifted LC ROI. B) For the rare xxxxx stimulus.

### 6. Comparison of pupil and fMRI responses to global deviants

Pupil response to auditory stimuli is controlled directly and indirectly by various neuromodulators (Joshi & Gold, 2020). We compared the responses evoked by global deviants in the subcortical ROIs (fMRI signal) and pupil size. We computed the average baseline-corrected pupil size in a window of 1.15-3 s after each global deviant (i.e. when pupil size showed an effect of global deviants at the group-level, see Figure 2), and splitted trials into large and small pupil responses (median split). Then, we compared the fMRI responses between those two conditions, expecting that deviant patterns accompanied by a larger pupil size would correspond to larger fMRI responses, in particular in the LC. Figure S4 shows the time course of the effect of global deviance (difference between frequent and rare patterns) in all sub-cortical ROIs conditioned on pupil response. This analysis is based on only 13 participants who had a large-enough number of trials per condition after artifact rejections (mean=67.6 trials, min=29, max=88); statistical power is thus low. The fMRI signal followed our prediction only in the SN/VTA (t_max_ = 4.05, p_max_ = 0.002, d_max_ = 1.12, cluster p_FWE_ = 0.022). No significant difference was found in the LC.

**Figure S5.**
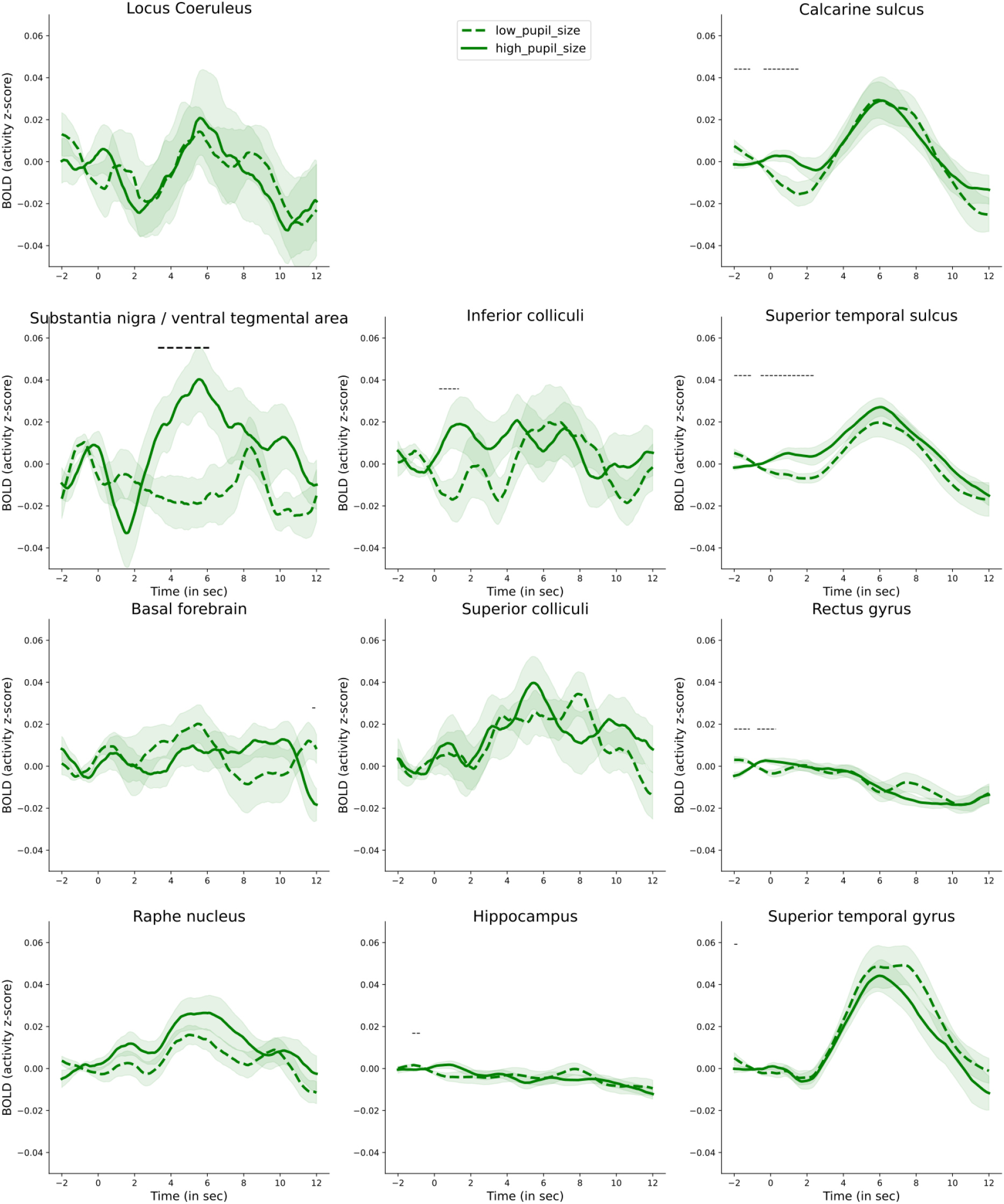
Time course of the effect of global deviance (z-scored fMRI activity elicited by rare), sorted by pupil response, in all subcortical ROIs. Horizontal dashed black lines correspond to clusters of significant difference (p < 0.05). Bold dashed black lines correspond to clusters of significant difference after FWE correction (p < 0.05).

### 7. Individual deliantion of the LC and the SN/VTA using TSE images

Figure S6 and Figure S7 present for each participant the slice in both TSE (one centered on the LC and one centered on the SN/VTA) for which each structure was the most visible.

**Figure S6.**
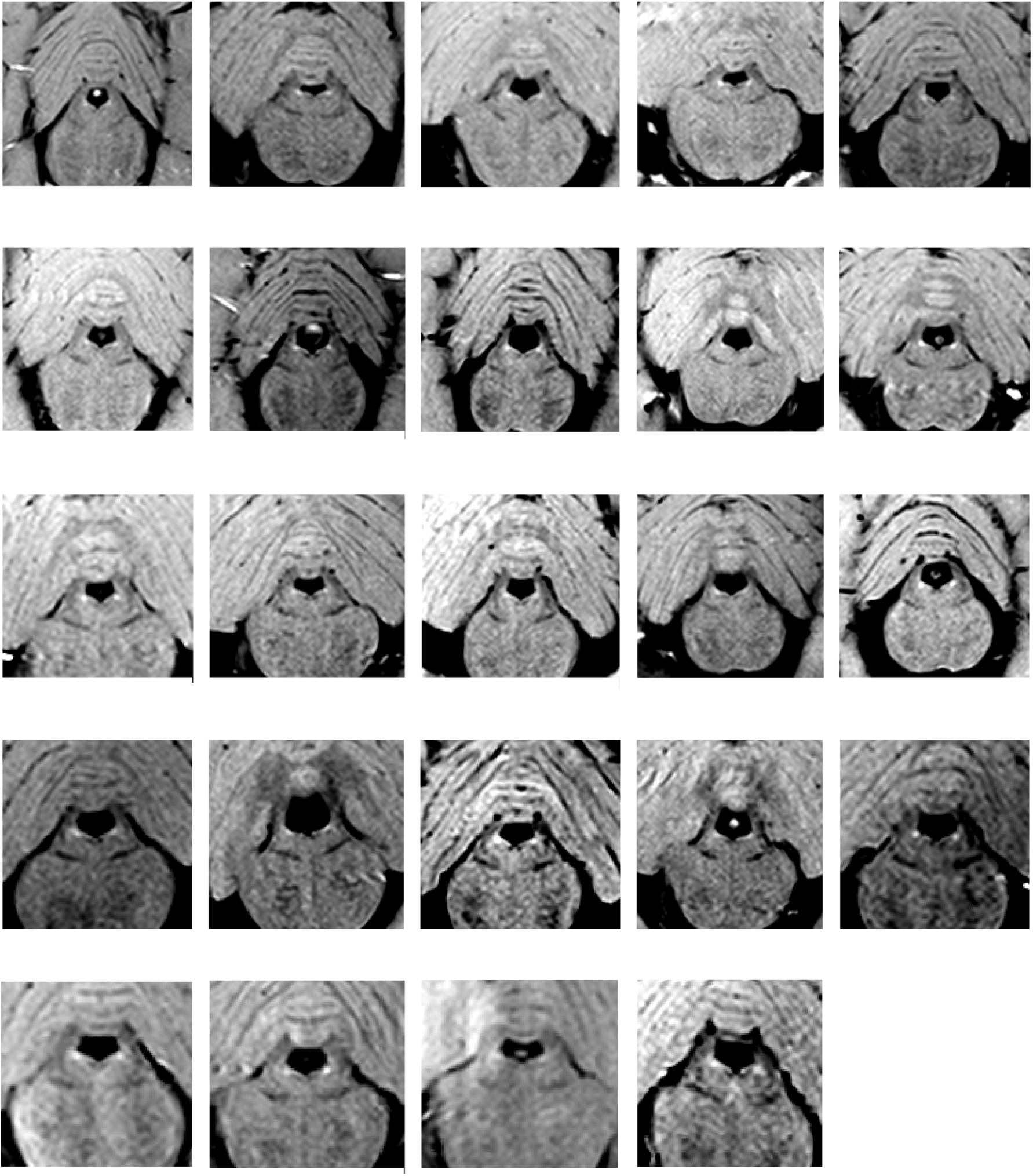
Anatomical images showing the LC in hypersignal for each participant.

**Figure S7.**
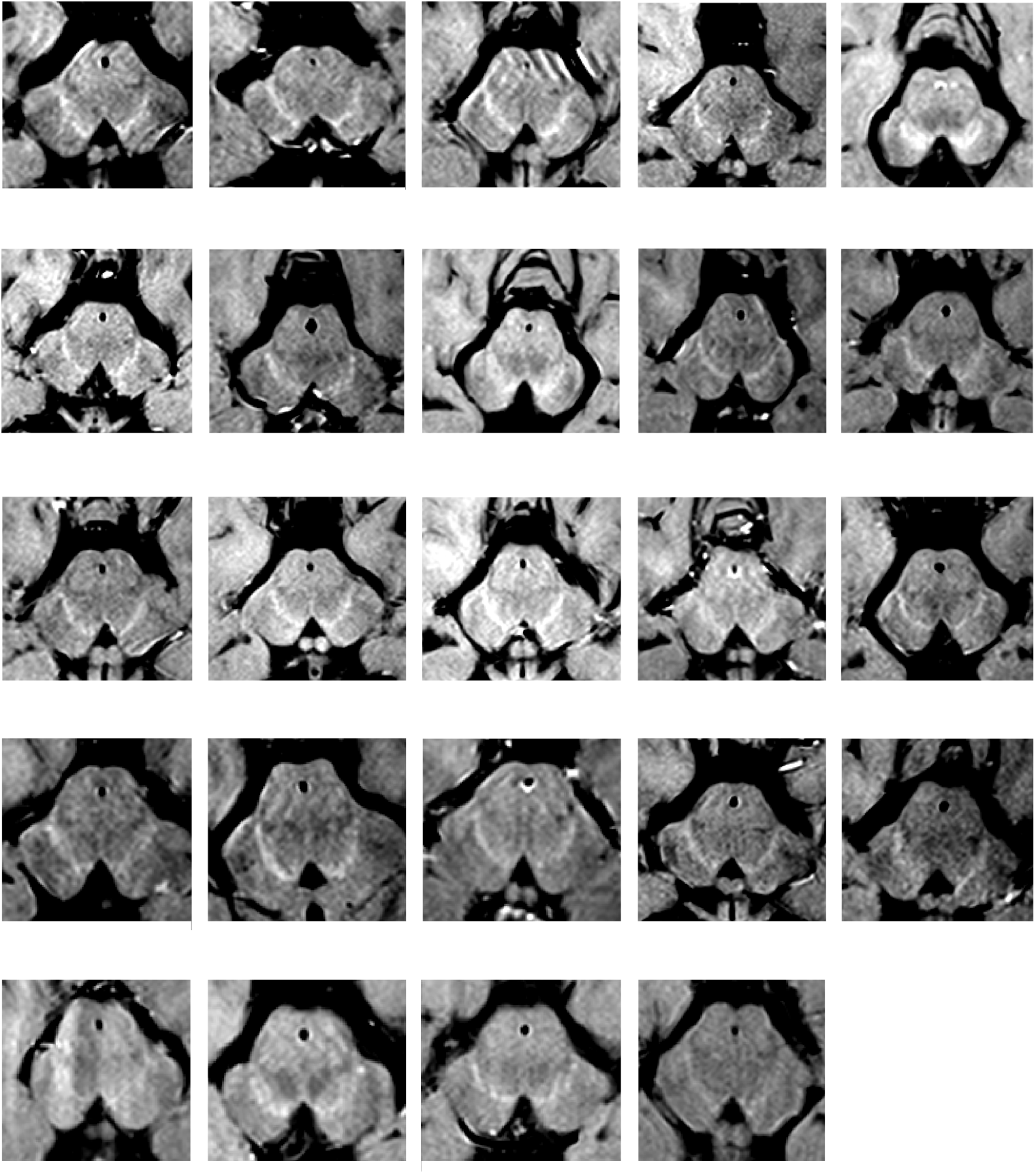
Anatomical images showing the SN/TVA in hypersignal for each participant.

## Notes

### Competing Interest Statement

The authors have declared no competing interest.

### Summary of Updates

- We have increased our sample size by 20%, rerun all analyses and redone all figures. The initial results are further confirmed with this larger sample size. - We have added an analysis of the correlation structures characterizing the subcortical regions of interest - We have clarified that the brain responses to rare patterns that we report cannot be uniquely attributed to a deviance detection process itself or to the consequences of this deviance detection in the task since rare patterns are task relevant. We have reworked the abstract and introduction to this end, and added a dedicated paragraph in the discussion. - We have added a number of additional analyses in the Supplementary Results

